# Intermediate-Resolution Modeling of Dynamic DNAs and Their Phase Separation

**DOI:** 10.64898/2026.09.14.751430

**Authors:** Jessica Fong Ng, Shanlong Li, Jianhan Chen

## Abstract

DNA is a fundamental biomolecule in eukaryotic cells, playing central roles in processes ranging from genome organization and transcription to innate immune signaling. Recent studies have revealed that DNA can undergo protein-free phase separation in the presence of divalent cations, yet the underlying molecular mechanisms, including the interplay of base stacking, base pairing, electrostatics, and ion interactions, remain poorly understood. Here, we introduce an intermediate-resolution model for condensates of DNAs (iConDNA) that can capture key local and long-range structural features of dynamic DNAs and simulate their spontaneous phase transitions. By introducing explicit base stacking and pairing interactions, the iConDNA model not only reproduces major conformational properties of DNA homopolymers but also folds DNA hairpins and duplexes and captures their thermodynamic properties. With an effective model of explicit Mg^2+^, iConDNA successfully captures the temperature and magnesium concentration dependence of DNA properties. Together, these features enable iConDNA to qualitatively recapitulate homotypic DNA phase separation, providing a suitable tool to study DNA homotypic phase separation in biological and engineering applications.

## Introduction

Biomolecule phase separation has emerged as a fundamental mechanism underlying cellular organization and processes. A growing body of evidence has established that phase separation drives diverse cellular processes, including transcription^1–4^, DNA repair, and stress response^5–7^. Mediated by multivalent interactions, spontaneous phase separation involves various biomacromolecules such as proteins and nucleic acids^8,9^. While intensive research interest has been drawn to condensates of proteins and/or RNA, DNA is not merely an observer, but an active participant in phase separation. Understanding the underlying mechanisms is therefore critical to elucidating its function across multiple cellular functions, from compartmentalization in chromatin at the nuclear level^10–13^, to the formation of condensates for stress response^14,15^, to immune responses triggered by cytoplasmic DNA^16,17^.

Recent studies have reported that DNA can undergo protein-free phase separation in the presence of divalent ions^18–21^. In particular, single-stranded DNA (ssDNA) undergoes thermos-responsive phase separation in the presence of divalent ions, such as Mg^2+^ or Ca^2+^, exhibiting lower critical solution temperature (LCST) behavior^22,18,21^. DNA trinucleotide repeats, including CAG_n_ and CUG_n_, can undergo Mg^2+^-mediated phase separation^21^. These DNA condensates strongly depend on Mg^2+^ concentration, temperature, sequence, and chain length, giving rise to complex phase-separation behaviors. However, the molecular mechanisms by which multivalent interactions and conformational dynamics collectively govern DNA phase separation remain poorly understood.

Molecular dynamics (MD) simulations have played an important role in elucidating the thermodynamic and mechanistic properties of DNA, as well as its folding and structural dynamics^13,23,24^. A range of coarse-grained (CG) models spanning different levels of resolution have been developed to make large-scale simulations tractable^25,26^. Minimal CG models can capture the thermodynamics of DNA hybridization^27–29^. Residue-based representations have been developed to reproduce the structural and mechanistic properties of B-form DNA^30–36^, while higher-resolution models can capture conformational transitions between A- and B-form DNA^37^. Beyond DNA-only systems, numerous CG models have been developed to study histone-DNA interactions and higher-order chromatin organization, providing insights into nucleosome positioning, chromatin fiber compaction, and loop extrusion^38,13,39,40^. More recently, the integration of machine learning has further expanded the accessible length and timescales of these studies^41–^ _43_ .

In this work, we developed an intermediate-resolution model for condensates of DNAs (iConDNA) by representing each nucleotide using six or seven CG beads and explicitly considering various backbone- and base-mediated interactions. Carefully calibrated DNA-specific backbone flexibility and geometries, along with interactions of base stacking and pairing, this model successfully reproduces experimentally observed structural and thermodynamic properties of ssDNA and double-stranded DNA (dsDNA). Furthermore, including an explicit representation of magnesium ions, iConDNA can capture the Mg^2+^-dependent structural properties and Mg^2+^-induced DNA phase separation. Taken together, iConDNA may provide a powerful tool for exploring the role of DNA in condensate-mediated biological functions and diseases.

## Methods

### Design of iConDNA

The iConDNA model maps each deoxyribonucleotide onto six or seven CG beads (**Fig. 1**). The phosphate group is represented by a single negatively charged bead (B1), which is connected to two beads representing the ribose sugar (B2 and B3). Each nucleobase is represented by a three- or four-bead ring for pyrimidines (cytosine, dC; thymine, dT) and purines (adenine, dA; guanine, dG), respectively. We adopted the same mapping scheme as in iConRNA^44^, except that thymine was modeled following the Martini DNA model^45^. As in the iConRNA model, B2 and B3 beads are fixed at the C4’ and C1’ positions of the ribose, respectively. The position of the remaining beads was determined from the corresponding centers of mass, except for the T2 bead of thymine, whose position was adjusted to reproduce the geometry of base pairing (**Supplementary Table 1**).

**Fig. 1.**
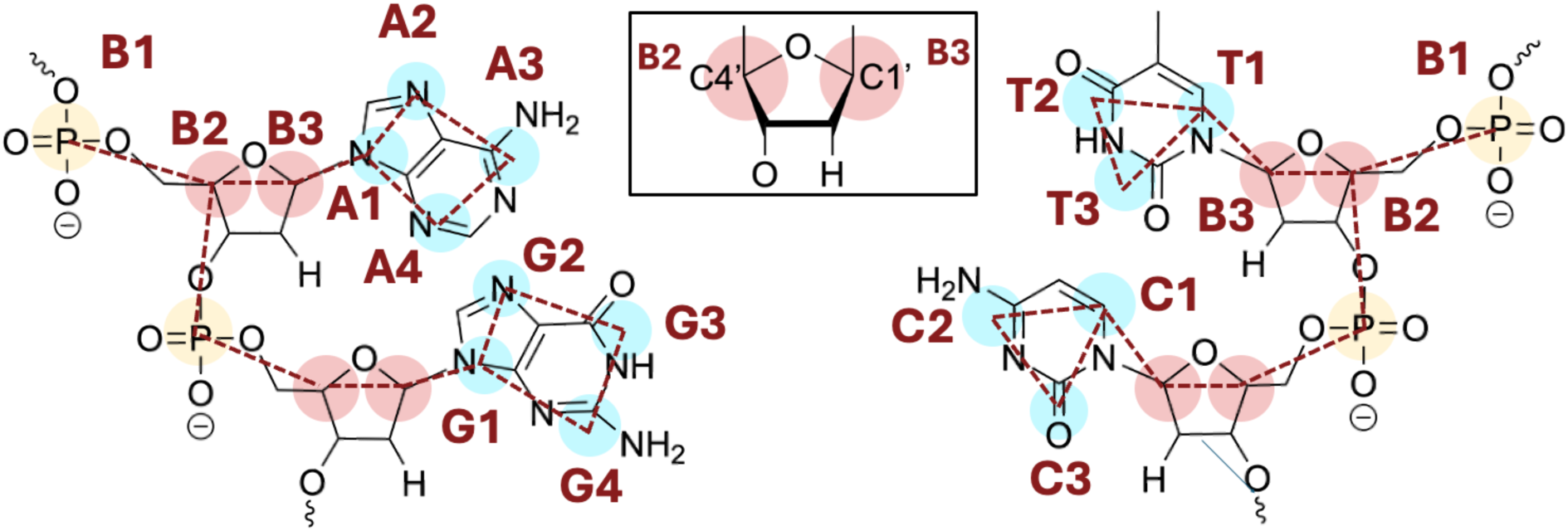
All-atom-to-CG mapping of DNA nucleotides in the iConDNA model. Colored circles represent CG beads for phosphate (yellow), sugar (red), and base (blue) groups. Bead names are labeled in red, while pseudo-bonds are indicated using red dashed lines. Sugar beads B2 and B3 correspond to the C4’ and C1’ atoms of the ribose, respectively, as shown in the inset.

### Energy function

The potential energy of this model includes four distinct contributions as previously described in our iConRNA model^44^

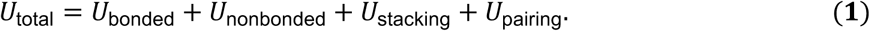

The *U*_bonded_ contains the bonded potentials for the intramolecular bonds, angles, dihedrals, and improper dihedrals. The *U*_nonbonded_ contains contributions from van der Waals (vdW) and Debye-Hückel-type electrostatic potential. *U*_stacking_ describes the stacking interactions among base beads between consecutive neighbors. Lastly, *U*_pairing_ is the base pairing potential between Watson-Crick base pairs. The details of each term are described in the following section.

Bonded Interactions. *U*_bonded_ contains standard bond, angle, dihedral, and improper dihedral terms, as used in previous iConRNA and HyRes protein models^46–48^

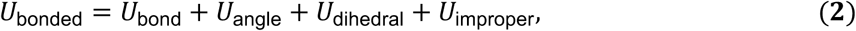

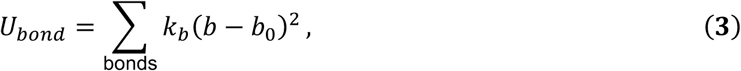

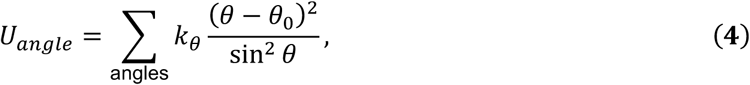

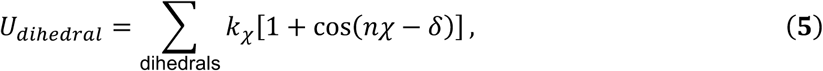

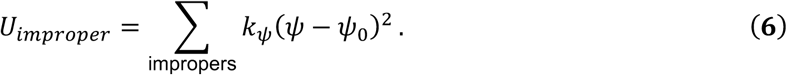

The parameters, including force constants and equilibrium values, were assigned to reproduce the distributions derived from all-atom (AA) explicit-solvent simulations of abasic trinucleotides. This molecule consists of an abasic sugar at the 5’ end, followed by one nucleotide and a sugar-phosphate backbone at the 3’ end (**Supplementary Fig. 2**). The absence of base stacking allows us to probe the intrinsic backbone dynamics.

Nonbonded Interactions. *U*_nonbonded_ includes the vdW and electrostatic terms

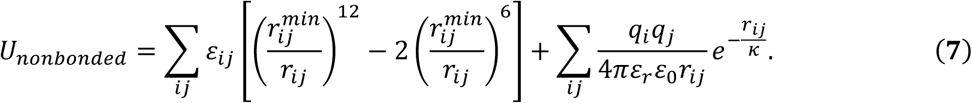

The distance between two beads *i* and 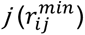 utilizes the Lorentz-Berthelot rules, where 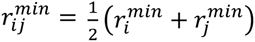. The interaction strength (ε_ij_) is given by 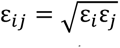. In the current version, only the backbone phosphate group (B1) carries a -1 charge, *q_B4_* = -1. ε% is the permittivity of vacuum. *κ* is the Debye screening length, determined as 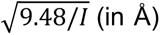, where *I* is the ionic strength in molarity.

The temperature dependence of electrostatic screening of water is captured by an empirical relation based on experimental dielectric constants (**Eq. 8**)^49^, where T represents the temperature in Celsius.

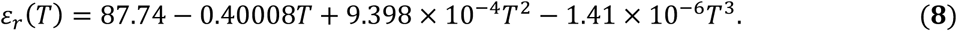

Since iConDNA sets *£_r_* = 60 at 300K following recent HyRes development^48^, thus this equation is scaled to

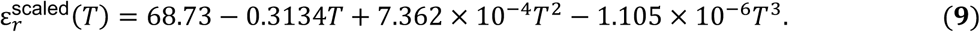

The initial vdW radius 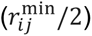 and interaction strength (*ε*) for each CG bead are taken from the same atom groups in the iConRNA force field^44^, or similar atom groups from the MARTINI force field^45^. The overall strength of vdW interactions is optimized by a single scaling factor during optimization to produce the structure and chain flexibility of poly(dT) and poly(dA) homopolymers, including radius of gyration (*R*_g_), end-to-end distance (*R*_e_), persistence length (*L*_p_), and orientation correlation function (OCF).

### Base stacking interactions

*U*_stacking_ is the stacking potential between neighboring bases (**Eq. 10**). The stacking interaction is applied only to consecutive bases from 5’ to 3’ using virtual sites to help mimic the B-form-like helical structure of DNA.

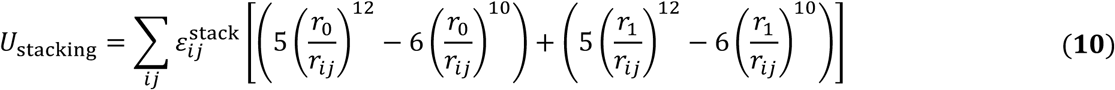

Two virtual bonds are applied between each base, with equilibrium distances *r_0_ and r_1,_* respectively, and 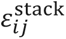 is the stacking strength between *i*and *j.* In the current model, the 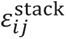 only depends on the types of stacking bases (denoted as *i*//*j*).

The virtual sites and bonds are designed based on the combination of purine (P, including dA and dG) and pyrimidine (R, including dC and dT), resulting in six (PP, PR, RR, RP) sets of defined interactions (**Supplementary Fig. 7** and **Supplementary Table 12**). The virtual sites and bonds are chosen manually based on the CG structure of 19 B-form duplexes from the PDB database (PDB IDs listed in **Supplementary Table 11**). The equilibrium stacking distances, *r_0_* and *r_1_*, reflect the calculated average from these structures (**Supplementary Table 7**).

The 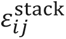 was determined by a relative strength table, where A//A is referenced as 1. The original table is taken from iConRNA and further adjusted according to experimental free energy calculations^50–53^ (**Supplementary Table 6**). Notably, T//T has stronger stacking interaction compared to U//U, and several base pairs are adjusted with higher strengths to reflect the experimental observation (A//T, C//C, C//T, G//C, G//T, T//C). The A//A stacking strength is calibrated with experimentally measured *R*_g_ values of dA_n_ and the structural propensity of dA_30_.

The structural propensity is quantified by the orientation correlation function (OCF)

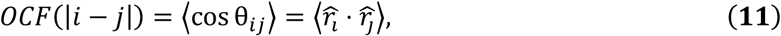

where 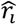 is the normalized bond vector between the *i_th_* and *(i + 1)_th_* phosphate groups along the DNA chain.

### Base pairing interactions

The canonical Watson-Crick base pairing is explicitly described in this model. For each base pair (*i* and *j*), two pseudo-hydrogen bonds are formed to mimic base pairing, where the pairing strength is controlled by:

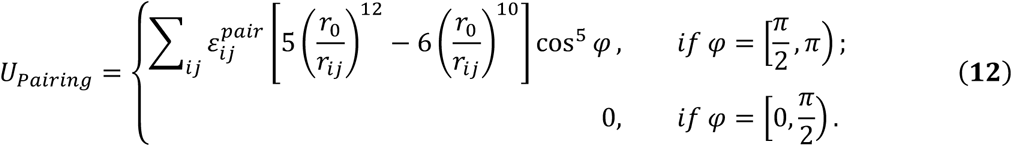

For an dA-dT pair, the pseudo-bonds are applied between beads A3-T2 and A4-T3, and for a dG-dC pair, they are applied between G3-C2 and G4-C3 (**Supplementary Fig. 9**). To maintain the planarity of the hydrogen bond, the position of T2 is adjusted from the center of mass of the mapped atoms to the center of mass of (N3, C4, O4) (**Supplementary Table 1**). The angles formed by A2-A3-T2 and G2-G3-C2 (ψ) are used to maintain the geometry of the hydrogen bond. The equilibrium distance *(r_0_)* and angles (ψ) are calculated as averages from PDB structures (**Supplementary Tables 8 and 9**).

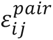 is the pairing strength, which was calibrated by reproducing the melting profiles of CTG repeats and the hairpins containing various GC content for G-C and A-T pairing, respectively (**Fig. 4A**, **Supplementary Tables 13** and **14**, and **Supplementary Fig. 10** and **11**). After that, the base pairing parameters were further validated using another set of hairpins and duplexes with various GC content (**Fig. 4B**, **Supplementary Table 16** and **17,** and **Supplementary Fig. 12** and **13**).

### Magnesium treatment and interactions

Homotypic phase separation of DNA requires the presence of crowding agents or divalent ions^18–21^. Although the screening of electrostatic interactions due to monovalent ions can be treated effectively using the Debye-Hückel-type electrostatic potential, this mean-field treatment is not sufficient for divalent ions because of ion-ion correlation and complex coordination with the phosphate backbone^54^. Mg^2+^ can interact with DNA through electrostatic interaction via a fully hydrated ion, or first-shell coordination that mediates polarization and charge transfer^55^. Their binding with DNA also depends on monovalent salt concentration, sequence, structure, and temperature^56–58^, and plays important roles in structural folding and stability^59^.

Following the iConRNA model, Mg^2+^ ion was explicitly described as a fully hydrated *[Mg(H_2_0)6]^2+^* (Mg^2+^) with a radius of 0.3 nm and a +2 charge^60^. The interaction between Mg and the phosphate group contains both vdW interaction (*ε* = 0.1 kcal/mol) and electrostatic interaction,

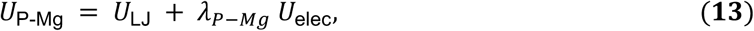

where λ_P-Mg_ was used to effectively describe the dependence of interaction strength on several conditions, including salt concentration, Mg^2+^ concentration, temperature, and RNA/DNA structure. In the iConRNA model, λ_P-Mg_ was finally fit against excess Mg^2+^ ions per phosphate (*Δn*_Mg_) by

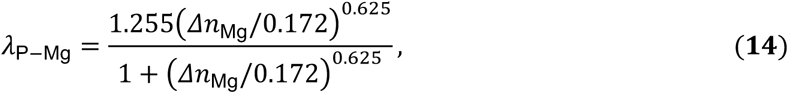

where *Δn*_Mg_ *= ΔN*_Mg_*/N*_P_. *ΔN*_Mg_ is the number of excess Mg^2+^ in the RNA-containing sample compared to the bulk buffer, and *N*_P_ is the number of phosphate groups^55,60–62^. *Δn*_Mg_ can be measured or predicted through atomistic simulations, theoretical models, or machine learning tools for various systems^55,60–65^. *ΔN*_Mg_ has been used to describe ssDNA properties with magnesium^65,66^. We would examine whether the relationship between *ΔN*_Mg_ and λ*_P-Mg_* via Eq. 14 is sufficient to describe DNA-Mg^2+^ interactions and capture the properties and phase separation propensity of ssDNA.

### Simulation details and analysis

#### Atomistic simulations

Atomistic simulations were performed using GROMACS 2026.3 package^67,68^ in the NPT ensemble under periodic boundary conditions. Abasic trinucleotides were constructed using PyMOL^69^. The AMBER99 force fields^70^ with the TIP3P water model^71^ were used for both DNA and RNA simulations. For DNA, the parmbsc1^72^ correction was applied, whereas the parmbsc0 and OL3^73,74^ corrections were applied for RNA. Initial systems were prepared using GROMACS. Long-range electrostatic interactions were calculated using the particle-mesh Ewald method^75,76^, and a cutoff of 1.1 nm was used for both short-range electrostatic and vdW interactions. Temperature was maintained using the V-rescale thermostat^77^ with a coupling time constant of 1.0 ps, while pressure was maintained at 1 bar using the Parrinello-Rahman barostat^78^ with a coupling time constant of 5.0 ps.

#### CG simulations

All CG simulations were performed using OpenMM 8.5.2^79^ in the NVT ensemble. Langevin dynamics were used with a friction coefficient of 0.1/ps. In-house Python packages were used to create the initial CG DNA or convert atomistic models to iConDNA representation.

#### Condensate simulations

The monomer structure was obtained from the AlphaFold^80^ server and converted to CG via in-house Python packages. The initial system was prepared using PACKMOL^81^ packages. Periodic boundary conditions were applied, and the system was equilibrated and run in the NVT ensemble using a Langevin middle integrator with a coupling time of 10ps.

#### Melting profile determination

Initial hairpin structures were obtained from the Protein Data Bank^82^ or predicted using the AlphaFold server^80^ and subsequently converted to CG structures using an in-house Python script. Simulations were performed at different temperatures for 6 μs for hairpins and 10 μs for duplexes. The first 50 ns and 200 ns of the hairpin and duplex trajectories, respectively, were discarded as equilibration periods prior to analysis. The number of intact base pairs was quantified from the trajectories using the MDAnalysis package^83,84^, and statistical convergence was assessed using block averaging. To determine the melting temperature, the average number of intact base pairs obtained from each block at each temperature was fitted to a sigmoidal curve using nonlinear least-squares fitting.

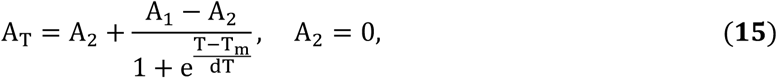

where *A_T_* represents the observed base pair at temperature *T*. *A_1_* and *A_2_* are the baseline values before and after the transition. *dT* is the slope factor describing the sharpness of the melting curve, and *T_m_* is the temperature at which half of the base pairs are observed. The reported standard deviation represents the variation in the melting temperatures obtained from the individual fits.

#### Sugar phase pseudorotation analysis of all-atom simulation

The MDAnalysis^83,84^ Nucleic Acid Analysis module is used to perform the phase angle analysis for atomistic abasic trinucleotides. The Cremer and Pople (CP) method^85^ is used to describe the geometry of the puckering relative to the plane by amplitude and phase coordinates.

## Results

### Parameterization and Validation

iConDNA followed a sequential parameterization, like iConRNA, where we combined both bottom-up and top-down approaches by using all-atom simulations, theoretical analysis, and experimental measurements. During the parameterization, we first established the bonded interactions, followed by nonbonded, then base-stacking, and lastly base-pairing interactions.

Abasic trinucleotides were used as reference molecules to generate atomistic distributions of bonded terms for characterizing DNA-specific backbone dynamics (**Supplementary Fig. 2**). The resulting CG distributions reproduced the overall profiles of the all-atom (AA) reference distributions, while smoothing out some of their finer features (**Supplementary Fig. 3-6**). Specifically, softer potentials were used for the B1-B2-B3 and B2-B1-B2* angles to encompass both the major and minor populations observed in the AA distributions.

DNA sugar puckering can alter the relative orientation of the phosphate backbone and nucleobase. We therefore examined how such conformational changes from the atomic simulations affect our CG bonded parameters. The atomistic simulations revealed that all four DNA nucleotides (dA, dT, dC, and dG) predominantly adopted the C2’-endo conformation, with less than 1% of the population adopting the C3’-endo conformation (**Supplementary Fig. 1A**). This distribution differs markedly from that of RNA counterparts, where both C2’- and C3’-endo conformations are substantially populated. Among the four RNA nucleotides, uracil showed the greatest flexibility, with approximately equal populations in the two conformational states (**Supplementary Fig. 1B**). Based on these DNA-specific conformational distributions, we expected our model to capture the distinct backbone conformational preferences of DNA relative to RNA (**Fig. 2**). Indeed, the CG backbone angles and dihedrals aligned well with atomistic distributions while remaining distinct from those of RNA (**Fig. 2A**). In addition, the orientation of the nucleobase relative to the ribose ring can adopt both syn and anti-conformations, as captured by the B2-B3-A1/G1-A2/G2 dihedral (**Fig. 2B**).

**Fig. 2.**
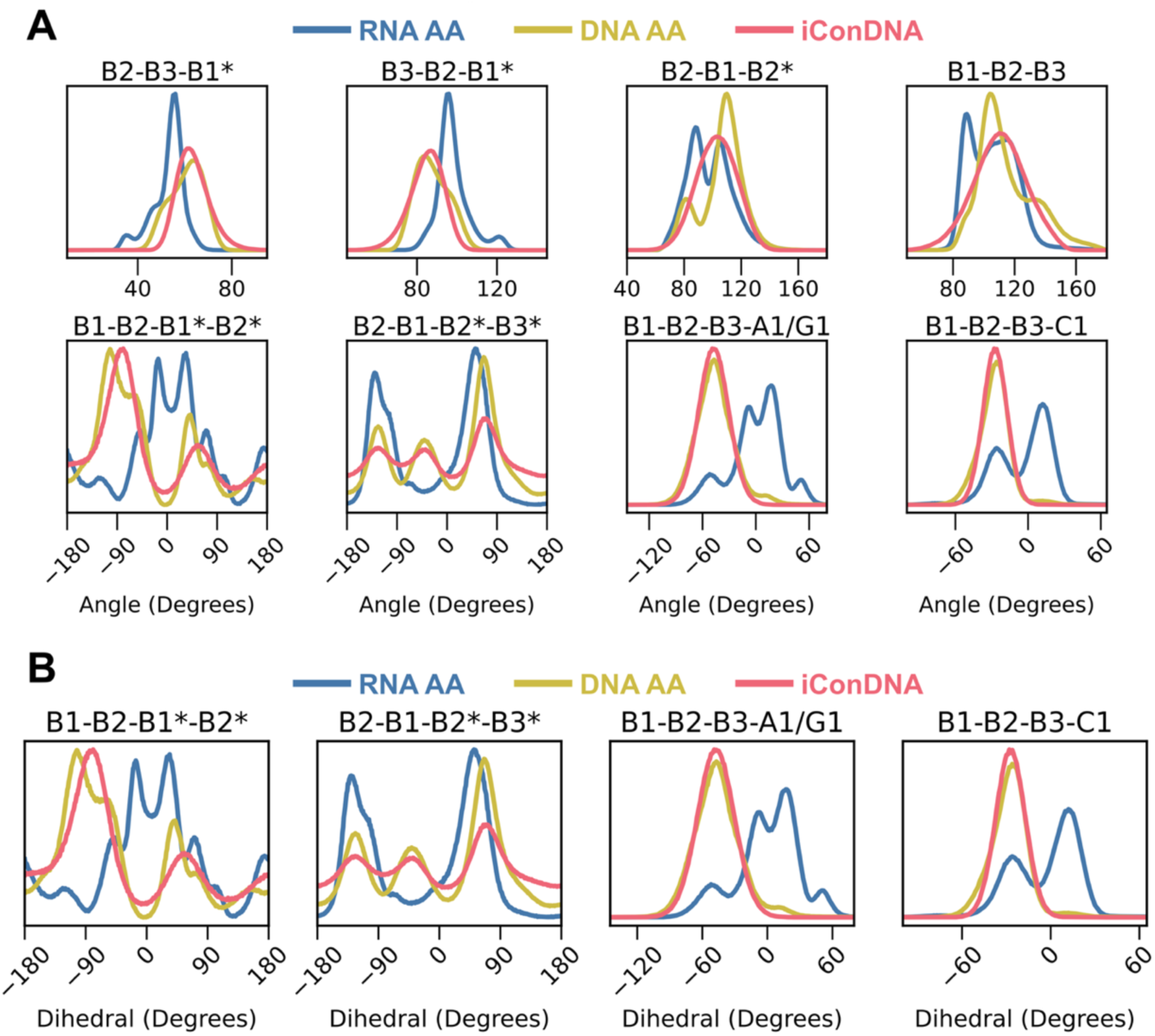
iConDNA accurately reproduces DNA-specific backbone geometry. Distributions of backbone angles and dihedrals from atomistic DNA (green), RNA (blue), and iConDNA (red) showing (**A**) phosphate-sugar backbone geometry and (**B**) base rotation of abasic trinucleotides. Asterisks (*) indicate beads from the next residue.

Overall, the dihedral force constants are weaker in this DNA model than those in the iConRNA model, resulting in a more flexible backbone^86–88^. Several angle force constants were increased to preserve sugar-ring geometry and maintain backbone stability. Specifically, the force constants of the B2-B1*-B3 and B2-B3-B1* angles were increased to 60 and 40 kcal/mol, respectively (**Supplementary Table 3**).

### Nonbonded, base-stacking, and base-pairing interactions

dT-dT has minimal base-stacking interaction^89,90,65^, making poly-dT an ideal reference system for parameterizing vdW interactions (**Fig. 3A** and **Supplementary Fig. 7).** Experimentally, dT_30_ adopts random coil conformations with a featureless OCF profile^66^, a feature well reproduced by our simulations (**Fig. 3A**). Although the model underestimated *R*_g_ and *L*_p_ relative to experiments^91,92^, it captured their salt-dependence, suggesting a reasonable balance between electrostatic and vdW interactions. The lower *R*_g_ and *L*_p_ suggest that the simulated DNA is overly flexible, whereas the higher *R*_e_ indicates excessive rigidity^91^. This apparent discrepancy may partly reflect differences between the simulated and experimentally measured observables. In particular, *R*_e_ was determined by FRET using fluorophore-labeled DNA, whereas the simulations contain only the DNA chain. The attached fluorophores and linkers, as well as the assumptions used to convert PRET efficiency to *R*_e_, may therefore contribute to differences between simulations and experiments. In addition, the experimentally observed non-monotonic dependence of *R*_g_ on salt concentration, including a decrease near 250 mM followed by an increase at 500 mM, was not reproduced by the CG simulations.

**Fig. 3.**
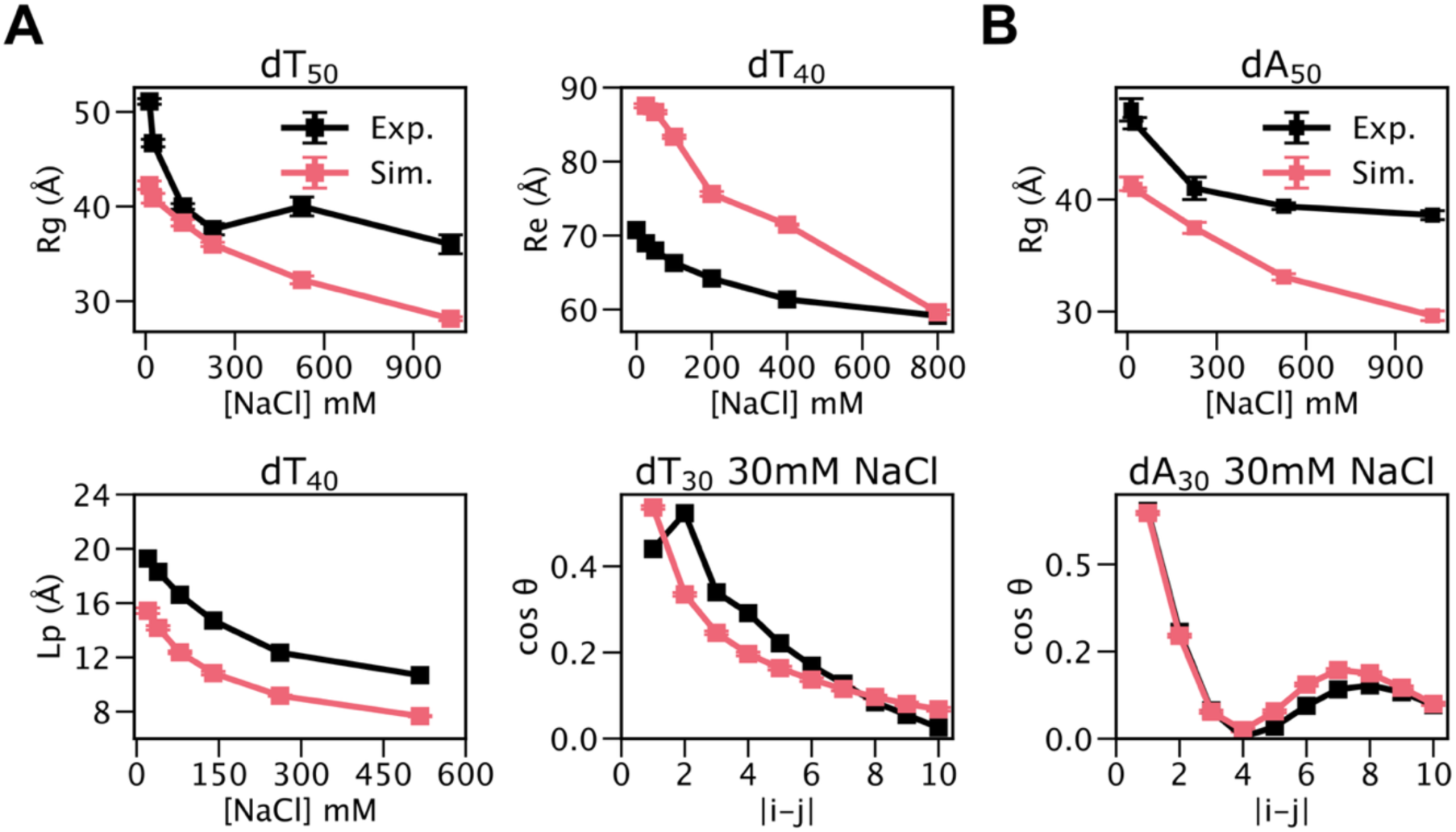
NaCl concentration-dependent dimensional and structural properties of DNA homopolymers. (**A**) NaCl concentration-dependent properties of poly(T)j, including the *R*_g_ of dT_50_ ^92^ (top left), *R*_e_ of dT_40_ ^91^(top right), *L*_p_ of dT_40_ ^91^(bottom left), and OCF of dT_30_ ^66^(bottom right). (**B**) NaCl concentration-dependent properties of poly(A), including *R*_g_ of dA_50_ (top) and OCF of dA_30_ (bottom). Experimental results are shown in black and simulation results in red. Error bars for simulation results were estimated using block averaging.

**Fig. 4.**
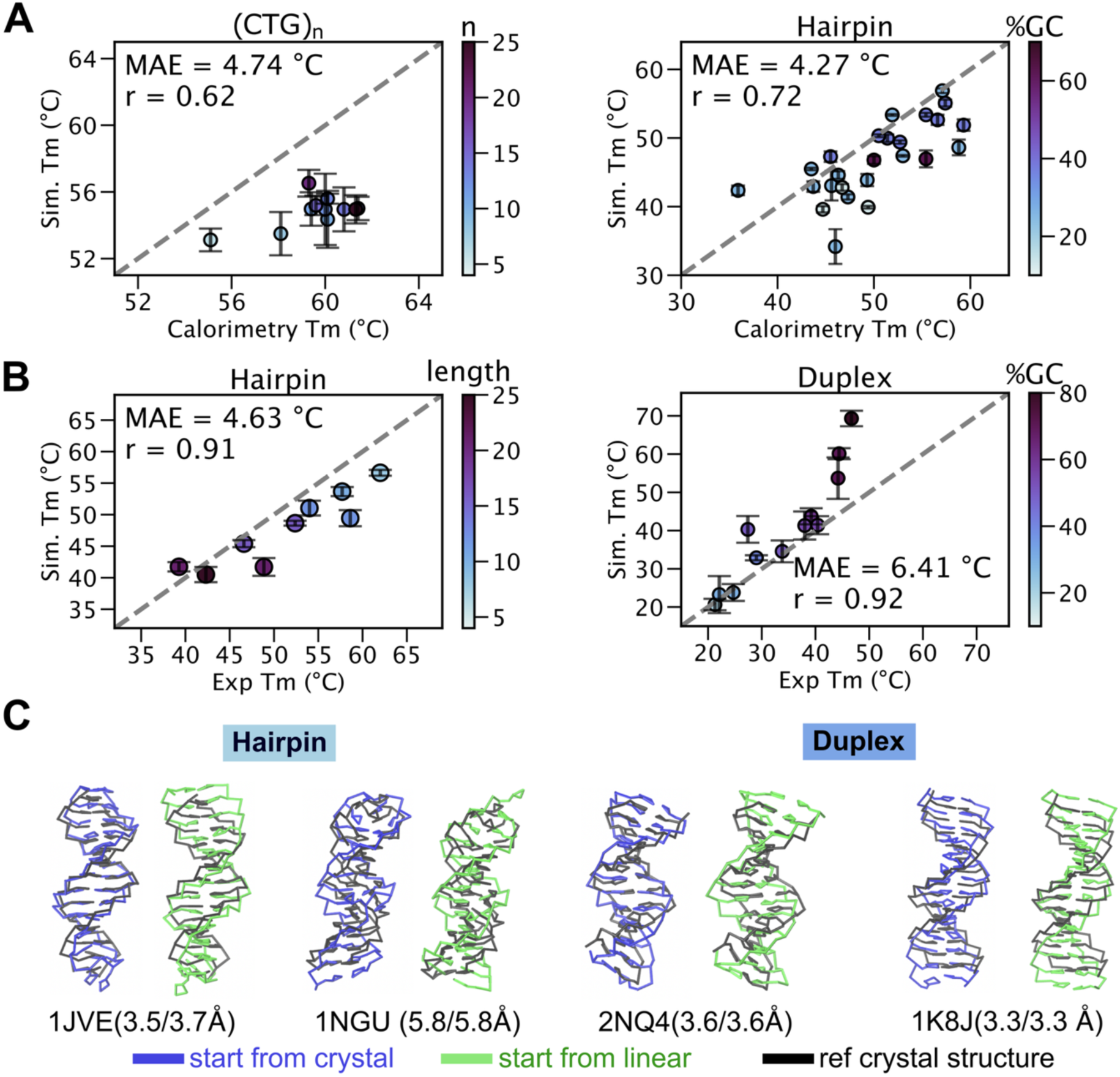
Melting profiles and folding of ssDNA and dsDNA. (**A, B**) Correlation between experimental and simulated melting profiles of hairpins and duplexes. (**A**) Base-pairing strength tuning sets using CTG_n_ (left) and hairpins (right). (**B**) Base-pairing testing sets using hairpins (left) and duplexes (right). Pearson correlation coefficients (r) and mean absolute errors (MAE) are shown in the upper left corners. (**C**) Comparison of experimental crystal structures (black) with predicted structures of ssDNA and dsDNA, starting from either crystal structures (blue) or linear chains (orange). These simulations were performed for 500 ns, with the first 100 ns discarded as equilibration.

Building on this baseline, we next parameterized base-stacking interactions using poly-dA data. The vdW radii were scaled down to reproduce the stacking geometry of B-form DNA. The final A//A stacking interaction was assigned a strength of 2 kcal/mol (eTDT’FG = 2 kcal/mol), yielding a total stacking strength of 4 kcal/mol between each pair of neighboring stacking interactions. With these parameters, the model successfully reproduced the helical structure of dA_30_, consistent with the experimentally reported OCF profile as well as *R*_e_ and *R*_g_ values^66,92^ (**Fig. 3B** and **Supplementary Fig. 7**).

Watson-Crick base-pairing interactions were optimized against experimentally measured melting temperatures. First, the G-C pairing strength was calibrated to reproduce the melting behavior of CTG repeats^93^, after which the A-T pairing strength was optimized using DNA hairpins with varying GC content^94^ (**Fig. 4A** and **Supplementary Figs. 10** and **11**; sequences are listed in **Supplementary Tables 12** and **13**). The model captured the length-independent melting profiles of CTG_n_ but failed to reproduce those of CAG repeats^93^ (**Supplementary Fig. 14**), suggesting that the stacking interactions involving dA require further refinement.

We next evaluated the base-pairing parameters using a set of hairpins containing various loop lengths^95^ and a set of duplexes with various GC content^96^ (**Fig. 4B**, **Supplementary Figs. 12** and **13**, and **sequences listed in Supplementary Tables 16 and 17**). Overall, the predicted melting temperatures of both hairpins and duplexes were in good agreement with experimental measurements. However, simulations of duplexes near the melting temperature were difficult to converge, with equilibrium not reached even after 10 µs of simulation (**Supplementary Fig.13**).

Finally, we evaluated the ability of the iConDNA model to fold small DNA structures. Starting from experimentally determined crystal structures or unstructured single strands, we simulated a series of simple DNA structures, including hairpins and duplexes (Fig. 4C). The model successfully formed B-form-like helices in both hairpin and duplex systems, although the predicted structures were not always consistent with the crystal structures.

### Magnesium-DNA interactions

The negatively charged phosphate backbone attracts cations from solution, resulting in an increased local cation concentration near the DNA surface relative to the bulk, a phenomenon known as counterion accumulation^54,97^. A corresponding decrease in the local dielectric constant has been observed experimentally and in simulations during DNA condensation^98–100^. To account for these changes in the local electrostatic environment, we reduced the dielectric constant from 60 to 20. This reduction strengthens phosphate-magnesium interactions, enhances the electrostatic repulsion between phosphate groups, and enhances the electrostatic screening by Mg^2+^.

The Mg^2+^ model followed the one used in the iConRNA model^44^. The interaction strength between phosphate and Mg is described by an empirical relationship between λ*_p-mg_* and Δ*n*_Mg_. We previously established this relationship using a Hill equation for RNA_44_ (**Eq. 10**) and tested whether it remains applicable to the DNA model. Using experimentally derived Δ*n*_Mg_ values^101^, the model reproduced the conformational properties of ssDNA dT_30_ across a range of Mg^2+^ concentrations (**Figs. 5A** and **B**). As discussed above, the discrepancy of underestimated *R*_e_ may partly arise from the absence of fluorescent dyes in the simulations, which are present in the FRET experiments used for comparison^102^.

**Figure 5.**
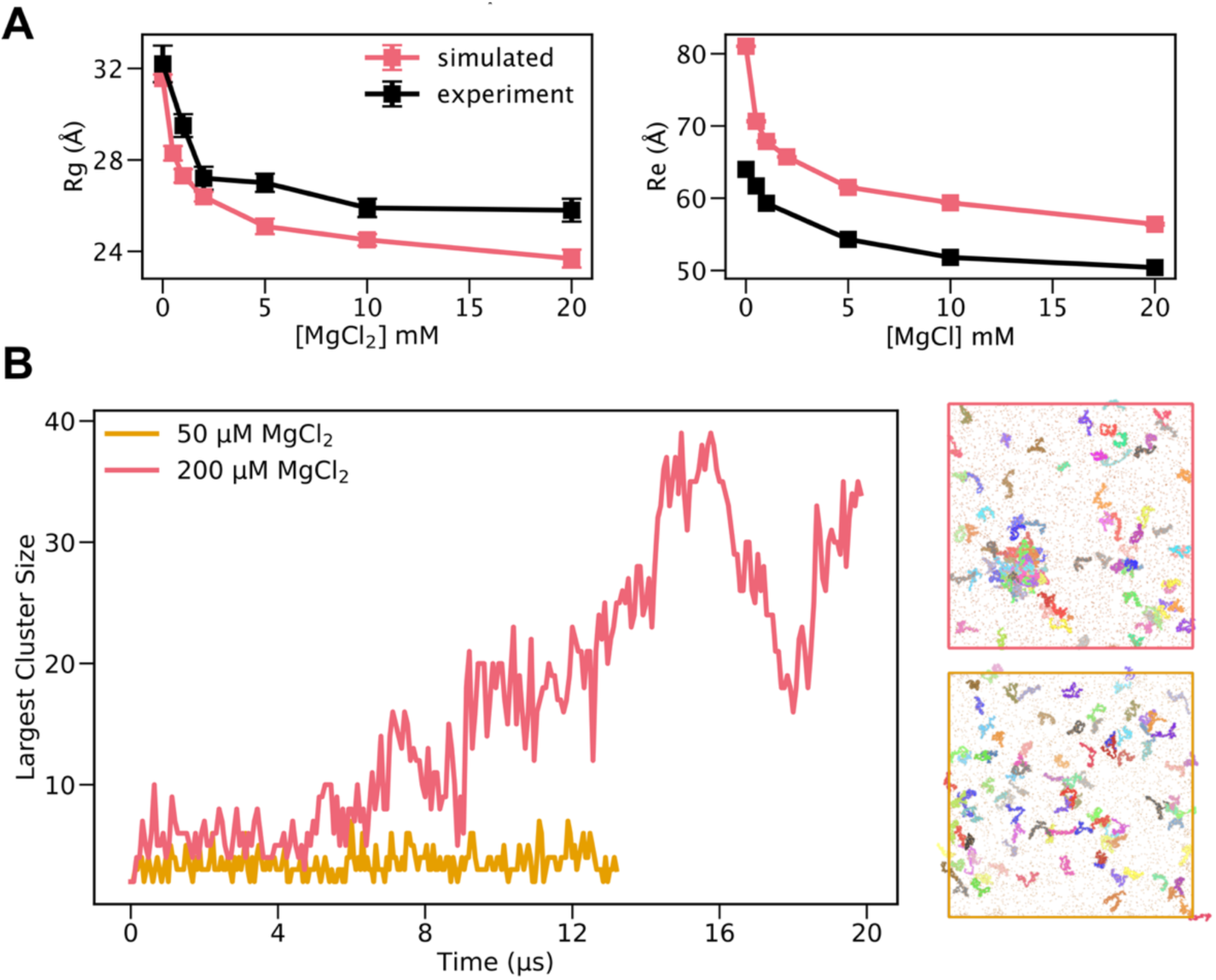
Mg^2+^-dependent ssDNA properties and Mg^2+^-mediated CAG_31_ phase separation. (**A**) Effect of MgCl_2_ concentration on dT_30_ R_g_ and R_e_ compared with experiment^102^. (**B**) Largest clusters formed across two simulated conditions: 100µM CAG_31_ with 50 mM MgCl_2_ at 80°C (yellow) and 100 µM CAG_31_ with 200 mM MgCl_2_ at 80°C (red). The representative snapshots from the end of the simulation are shown on the right.

### Mg^2+^-mediated DNA phase separation

We next examined whether the model could capture DNA phase separation using CAG_31_ (**Table 1**). At the experimentally observed condition of 100 µM CAG_31_ and 50 µM Mg^2+^ at 80 °C^21^, the model did not reproduce phase separation (**Fig. 5B**). Increasing the Mg^2+^ concentration to 200µM resulted in cluster formation (**Fig. 5B**), although the system remained highly dynamic and did not reach convergence even after 20 µs of simulation (**Fig. 5B**). This suggested that the propensity for DNA phase separation was underestimated in the iConDNA model.

**Table 1.** CAG_31_ simulated result compared to experimental observation.

| Condition | Phase Separation Propensity |  |
| --- | --- | --- |
|  | Experiment | Simulation |
| 50μM CAG<sub>31</sub> 80 °C | Yes | No |
| 50μM CAG<sub>31</sub> 50 °C | No | No |
| 100μM CAG<sub>31</sub> 80 °C | Yes | Yes |
| 100μM CAG<sub>31</sub> 65 °C | Yes | No |

We hypothesized that the vdW and stacking interactions parameterized at a dielectric constant of 60 may be insufficient to compensate for the increased electrostatic repulsion at a dielectric constant of 20. To test whether changing the dielectric constant substantially alters the intrinsic chain properties, we compare the *R*_e_ and *R*_g_ of dT_30_ at a dielectric constant of 20 and 60. The chain compaction was similar under both conditions (**Supplementary Fig. 15**), suggesting that the change in dielectric constant does not substantially alter the single-chain conformational properties. These results instead suggest that the current stacking interactions may be too weak to promote the intermolecular base-base interactions required for phase separation, consistent with the importance of such interactions in the iConRNA model.

## Discussion

In this work, we introduced iConDNA, an intermediate-resolution coarse-grained (CG) model designed to bridge molecular-level chemical detail with tractable sampling of large-scale DNA conformational dynamics and phase separation. By representing each nucleotide with six or seven CG beads and deriving bonded interactions from atomistic distributions of abasic trinucleotides, iConDNA directly captures the distinct conformational preferences of the DNA backbone. Notably, the model reflects the dominant C2′-endo sugar-puckering geometry that distinguishes DNA from RNA, avoiding excessive rigidity while maintaining backbone stability. Combined with carefully tuned base-stacking and canonical base-pairing interactions, iConDNA reproduces the ionic strength-dependent dimensions of homopolymers, successfully recapitulates the thermal stability of various DNA hairpins and duplexes, and spontaneously folds unstructured single strands into native-like B-form conformations.

A central objective of iConDNA is to elucidate the physical forces governing protein-free, divalent cation-mediated DNA phase separation. Simulating trinucleotide repeats demonstrated that the model qualitatively captures the lower critical solution temperature (LCST)- like behavior of single-stranded DNA, forming large dynamic clusters for CAG_31_ at elevated temperatures of 80 °C. However, the model underestimates the overall propensity for phase separation: clustering required an elevated DNA concentration of 100 µM rather than the experimentally observed 50 µM and failed to assemble at 65 °C.

Moving forward, several targeted refinements will help elevate iConDNA from qualitative consistency to quantitative predictive power. Parameterizing λ_P-Mg_ directly against DNA-specific ion-counting data will ensure accurate ion condensation across broader salt and temperature regimes. Expanding the potential functions to accommodate non-canonical base interactions, such as G–G pairings found in G-quadruplexes^103–105^, will broaden the model’s scope to biologically critical non-B-form structures. Finally, simultaneously co-optimizing van der Waals, stacking, and pairing parameters under a reduced dielectric regime will properly balance attractive dispersion forces against electrostatics in crowded environments. In its present form, iConDNA provides an efficient, physics-based platform for investigating DNA mechanics and phase transitions, establishing a solid foundation for multiscale studies of chromatin organization and synthetic DNA nanotechnology.

## Data, Materials and Software Availability

The data underlying this article, including all iConDNA force field files, are available from GitHub at https://github.com/jessica-fongng/iConDNA. All other data are included in the manuscript or SI.

## Acknowledgements

This work is supported by NSF through CHE 2516941. The computational resources for this work were provided by the University of Massachusetts Amherst’s partnership with the Unity Research Computing Platform, a multi-institutional cluster led by the University of Massachusetts and the University of Rhode Island.

## Author contributions

Chen conceptualized the idea. Fong Ng conducted the iConDNA development and performed simulations and analyses. Chen and Li contributed to the analysis. Fong Ng drafted the manuscript. Chen and Li revised the manuscript.

## Additional information

Extended data is available for this paper at https://doi.org/xxxx

## Supplementary information

The online version contains supplementary material available at https://doi.org/xxxx

## Supplementary Information

**Supplementary Table 1.**
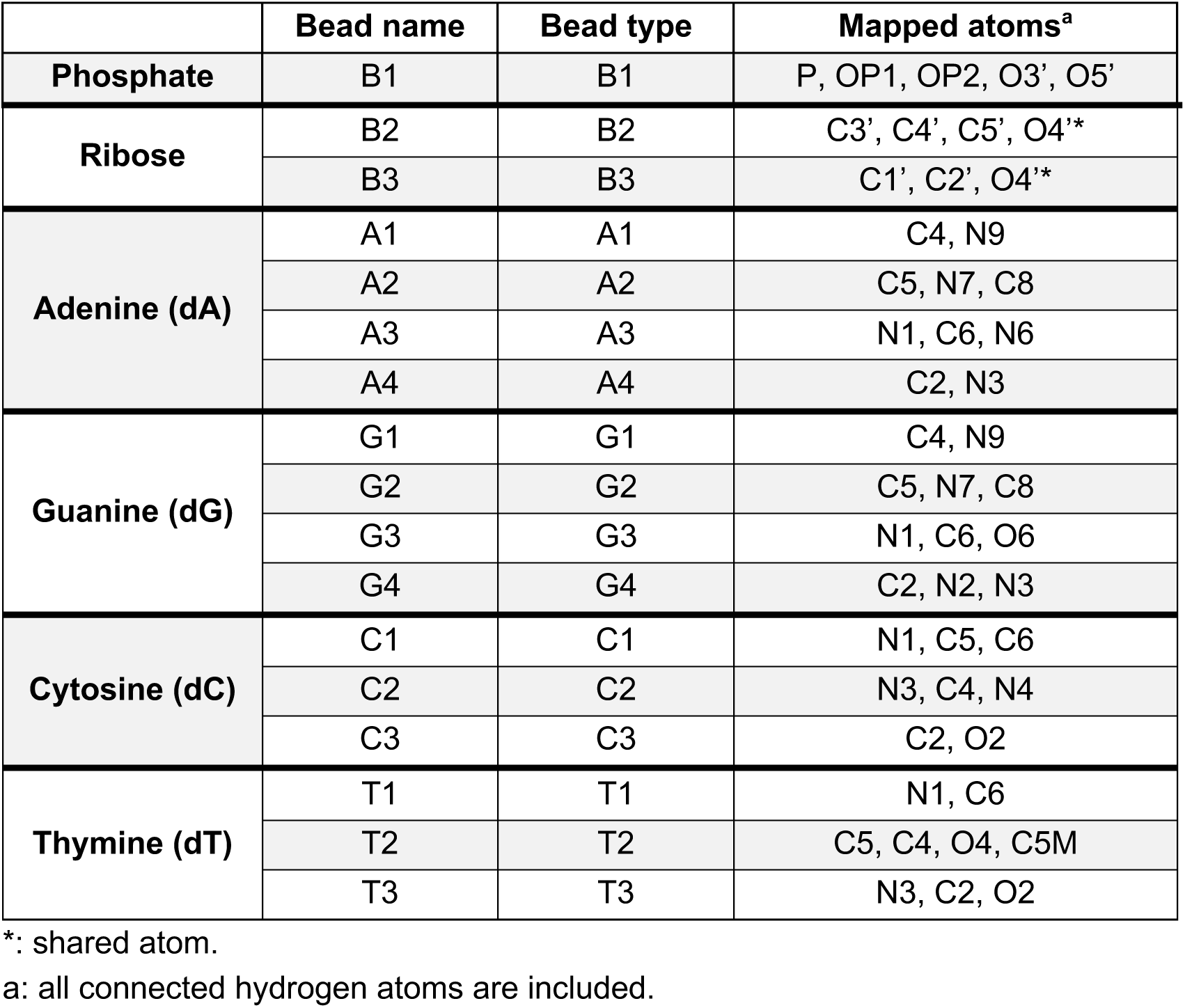
Atomistic to CG mapping scheme (also see Fig. 1).

|  | Bead name | Bead type | Mapped atoms<sup>a</sup> |
| --- | --- | --- | --- |
| **Phosphate** | B1 | B1 | P, OP1, OP2, O3', O5' |
| **Ribose** | B2 | B2 | C3', C4', C5', O4'* |
|  | B3 | B3 | C1', C2', O4'* |
| **Adenine (dA)** | A1 | A1 | C4, N9 |
|  | A2 | A2 | C5, N7, C8 |
|  | A3 | A3 | N1, C6, N6 |
|  | A4 | A4 | C2, N3 |
| **Guanine (dG)** | G1 | G1 | C4, N9 |
|  | G2 | G2 | C5, N7, C8 |
|  | G3 | G3 | N1, C6, O6 |
|  | G4 | G4 | C2, N2, N3 |
| **Cytosine (dC)** | C1 | C1 | N1, C5, C6 |
|  | C2 | C2 | N3, C4, N4 |
|  | C3 | C3 | C2, O2 |
| **Thymine (dT)** | T1 | T1 | N1, C6 |
|  | T2 | T2 | C5, C4, O4, C5M |
|  | T3 | T3 | N3, C2, O2 |

\*: shared atom.
a: all connected hydrogen atoms are included.

**Supplementary Table 2.** Bond parameters, *k_b_* in kcal/mol•Å and *b_0_* in Å.

| i | j | $k_b$ | $b_0$ | i | j | $k_b$ | $b_0$ |
| --- | --- | --- | --- | --- | --- | --- | --- |
| B1 | B2 | 30 | 3.94 | G1 | G4 | 150 | 2.97 |
| B2 | B3 | 100 | 2.32 | G2 | G3 | 125 | 2.97 |
| B3 | A1/G1 | 200 | 1.93 | G3 | G4 | 160 | 2.89 |
| B3 | C1 | 160 | 2.48 | C1 | C2 | 350 | 2.41 |
| B3 | T1 | 250 | 1.9 | C2 | C3 | 150 | 2.88 |
| A1/G1 | A2/G2 | 250 | 1.63 | C1 | C3 | 250 | 2.63 |
| A1 | A4 | 150 | 2.35 | T1 | T2 | 300 | 2.89 |
| A2 | A3 | 125 | 3.00 | T2 | T3 | 270 | 2.41 |
| A3 | A4 | 300 | 2.63 | T1 | T3 | 350 | 2.25 |

**Supplementary Table 3.** Angle parameters, *k_θ_* in kcal/(molᐧra.^2^) and *θ_0_* in degrees.

| i | j | k | $k_\theta$ | $\theta_0$ | i | j | k | $k_\theta$ | $\theta_0$ |
| --- | --- | --- | --- | --- | --- | --- | --- | --- | --- |
| B2 | B1 | B2 | 2 | 105 | B3 | A1 | A4 | 65 | 117 |
| B1 | B2 | B1 | 5 | 121.5 | B3 | G1 | G4 | 45 | 117 |
| B1 | B2 | B3 | 1 | 125.0 | B3 | C1 | C3 | 75 | 61 |
| B2 | B3 | B1* | 7<sup>†</sup> | 57.0 | B3 | T1 | T3 | 75 | 87.8 |
| B2 | B1* | B3 | 40<sup>‡</sup> | 32.0 | A1 | A2 | A3 | 50 | 90 |
| B2 | B3 | A1/G1 | 4 | 147 | A2 | A3 | A4 | 50 | 72 |
| B2 | B3 | C1 | 20 | 123 | A3 | A4 | A1 | 50 | 86.5 |
| B2 | B3 | T1 | 20 | 123.9 | G1 | G2 | G3 | 75 | 92.0 |
| B3 | A1/G1 | A2/G2 | 60 | 129.9 | G2 | G3 | G4 | 50 | 83 |
|  |  |  |  |  | G3 | G4 | G1 | 50 | 72.5 |

\* The bead in the next residue.
† Force constant set to 40 kcal/mol in the final model to improve stability; see main text.
‡ Force constant increased to 60 kcal/mol in the final model to improve stability; see main text.

**Supplementary Table 4.** Dihedral parameters, *kx* in kcal/(mol) and *S* in degrees.

| i | j | k | i | $k_\chi$ | n | $\delta$ |
| --- | --- | --- | --- | --- | --- | --- |
| B1 | B2 | B1 | B2 | 0.26 | 1 | 90.62 |
| B1 | B2 | B1 | B2 | 0.26 | 2 | -27.02 |
| B1 | B2 | B1 | B2 | 0.2 | 3 | -37.49 |
| B2 | B1 | B2 | B3 | 0.15 | 1 | -167.41 |
| B2 | B1 | B2 | B3 | 0.31 | 2 | -28.30 |
| B2 | B1 | B2 | B3 | 0.19 | 3 | 80.62 |
| B2 | B1 | B2 | B3 | 0.19 | 4 | 43.9 |
| B1 | B2 | B3 | A1/G1 | 7.0 | 1 | 132.3 |
| B1 | B2 | B3 | C1 | 20.0 | 1 | 152.8 |
| B1 | B2 | B3 | T1 | 20.0 | 1 | 154.2 |
| B2 | B3 | A1/G1 | A2/G2 | 0.77 | 1 | -160.7 |
| B2 | B3 | A1/G1 | A2/G2 | 126 | 2 | -65.0 |
| B2 | B3 | C1 | C3 | 10 | 1 | 16.81 |
| B2 | B3 | T1 | T3 | 10 | 1 | 20.56 |
| A1 | A2 | A4 | A3 | 80 | 1 | 0 |
| A1 | A2 | A3 | A4 | 50 | 1 | 180 |
| G1 | G2 | G4 | G3 | 50 | 1 | 0 |
| G1 | G2 | G3 | G4 | 80 | 1 | 180 |

**Supplementary Table 5.** Improper parameters: *k_ѱ_* in kcal/(mol) and *Ѱ_0_* in degrees.

| i | j | k | l | $k_\psi$ | $\Psi_0$ |
| --- | --- | --- | --- | --- | --- |
| B2 | B1 | B1* | B3 | 40 | 41.5 |
| A1 | B3 | A2 | A4 | 40 | 0 |
| G1 | B3 | G2 | G4 | 30 | 0 |
| C1 | B3 | C2 | C3 | 7.0<sup>†</sup> | -180 |
| T1 | B3 | T3 | T2 | 20.0 | 0 |

\* The bead in the next residue.
† Force constants set to 20kcal/mol to improve local structure.

**Supplementary Table 6.** Relative stacking strength to A//A stacking. In the final model, *ε_A//A_* = 4.0 kcal/mol.

|  | A//A | A//C | A//G | A//T | G//A | G//C | G//G | G//T |
| --- | --- | --- | --- | --- | --- | --- | --- | --- |
| $\epsilon_{\text{relative}}$ | 1 | 0.8 | 1 | 1 | 1 | 1 | 1 | 1 |
|  | T//A | T//C | T//G | T//T | C//A | C//C | C//G | C//T |
| $\epsilon_{\text{relative}}$ | 0.4 | 0.6 | 0.4 | 0.4 | 0.4 | 0.4 | 0.4 | 0.4 |

**Supplementary Table 7.** Equilibrium distance in Å for different stacking base pairs.

| Bond #1<br>($r_0$) | A//A | A//C | A//G | A//T | G//A | G//C | G//G | G//T |
| --- | --- | --- | --- | --- | --- | --- | --- | --- |
| Å | 3.5 | 3.6 | 3.8 | 3.4 | 3.5 | 3.7 | 3.7 | 3.5 |
| Bond #2<br>($r_1$) | A//A | A//C | A//G | A//T | G//A | G//C | G//G | G//T |
| Å | 4.4 | 3.6 | 4.1 | 3.4 | 4.4 | 3.4 | 4.2 | 3.3 |
| Bond #1<br>($r_0$) | T//A | T//C | T//G | T//T | C//A | C//C | C//G | C//T |
| Å | 3.6 | 3.6 | 3.6 | 3.7 | 3.7 | 3.6 | 3.8 | 3.6 |
| Bond #2<br>($r_1$) | T//A | T//C | T//G | T//T | C//A | C//C | C//G | C//T |
| Å | 4.1 | 3.6 | 4.1 | 3.6 | 4 | 3.6 | 4 | 3.7 |

**Supplementary Table 8.** Equilibrium distance for different hydrogen-bonding CG beads and optimized hydrogen-bonding pair strength.

| Base pair | Bead pair | $r_0$ (nm) | $\epsilon_{\text{pair}_{ij}}$ (kcal/mol) |
| --- | --- | --- | --- |
| A-T | A3-T2 | 0.327 | 2.78 |
|  | A4-T3 | 0.43 |  |
| G-C | C2-G3 | 0.33 | 4.17 |
|  | C3-G4 | 0.4 |  |

**Supplementary Table 9.** Equilibrium angle for different hydrogen-bonding CG beads and optimized hydrogen-bonding pair strength.

| Base pair | Bead pair | Angle $\theta_0$ (°) |
| --- | --- | --- |
| A-T | A2-A3-T2 | 180 |
| G-C | G2-G3-C2 | 180 |

**Supplementary Table 10.** NBFIX for vdW parameters.

| i | j | $\epsilon_{ij}$ | $r_{\text{min}}/2(\text{\AA})$ |
| --- | --- | --- | --- |
| B1 | MG | 0.500* | 5.0* |
| C2 | G3 | -0.12669 | 3.31 |
| C3 | G4 | -0.10337 | 4.01 |
| A3 | T2 | -0.12669 | 3.27 |

\*The parameters are adopted from iConRNA.

**Supplementary Table 11.**
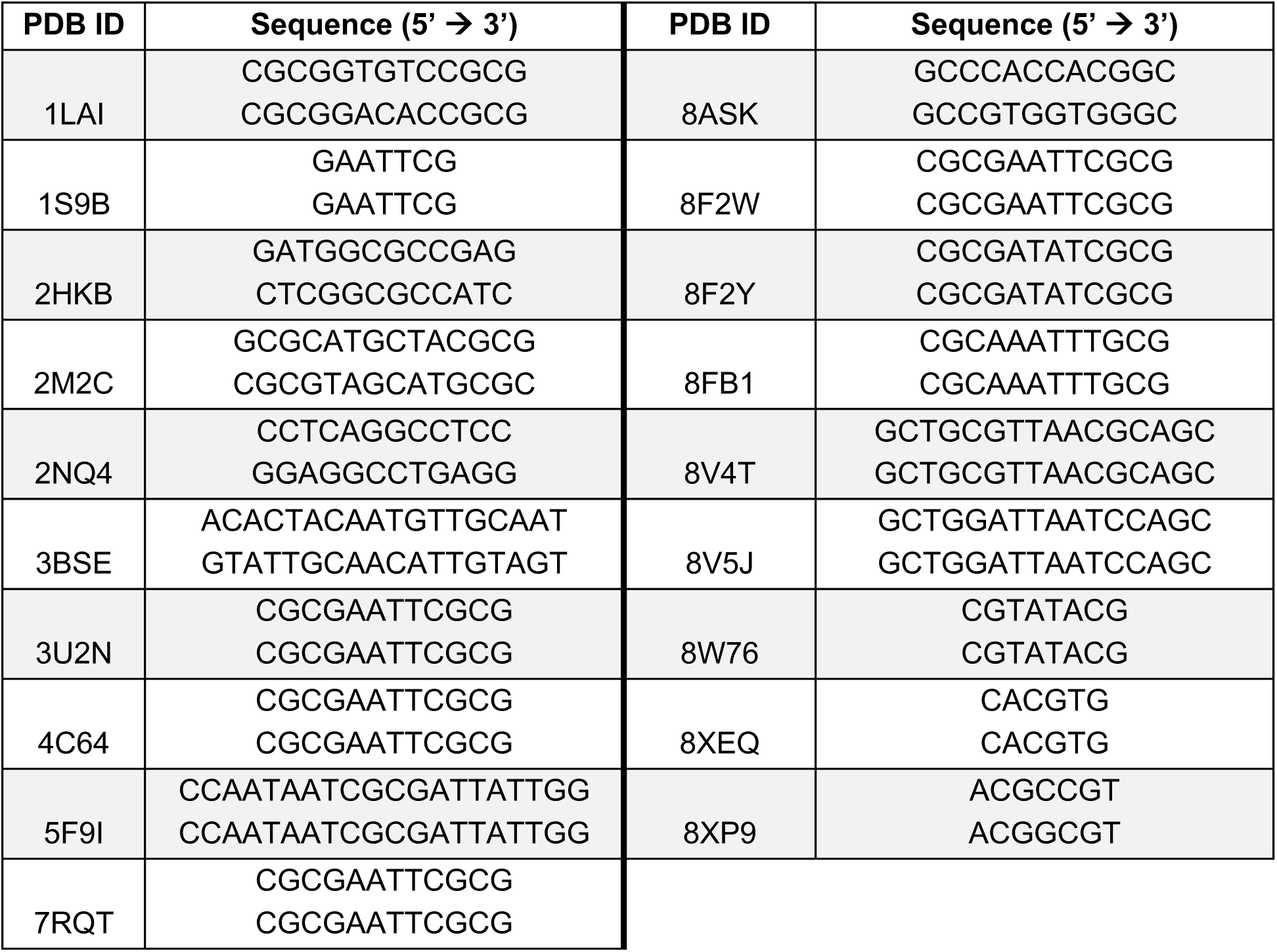
PDB used for base stacking and base pairing parameterization.

**Supplementary Table 12.** Bead(s) used for calculating the center of geometry as virtual sites for base stacking. Purines (R) correspond to adenine (A) and guanine (G). Pyrimidine (Y) refers to cytosine (C) and thymine (T).

| 5' → 3' | Bond # 1 |  | Bond #2 |  |
| --- | --- | --- | --- | --- |
|  | Virtual site 1 | Virtual site 2 | Virtual site 3 | Virtual site 4 |
| R to R | A4/<br>G4 | A1, A2, A3, A4/<br>G1, G2, G3, G4* | A2/<br>G2 | A1, A2, A3, A4/<br>G1, G2, G3, G4* |
| R to Y | A1, A2, A3, A4/<br>G1, G2, G3, G4 | C1, C2/<br>T1, T2 | A4/<br>G4 | C3/<br>T3 |
| Y to Y | C3/<br>T3 | C1, C2, C3/<br>T1, T2, T3<sup>†</sup> | C1, C2, C3/<br>T1, T2, T3<sup>†</sup> | C2/<br>T2 |
| Y to R | C3/<br>T3 | A2, A3/<br>T2, T3 | C3/<br>T3 | A1, A4/<br>T1, T4 |

\*†‡The virtual site is shared between bond #1 and bond #2.

**Supplementary Table 13.**
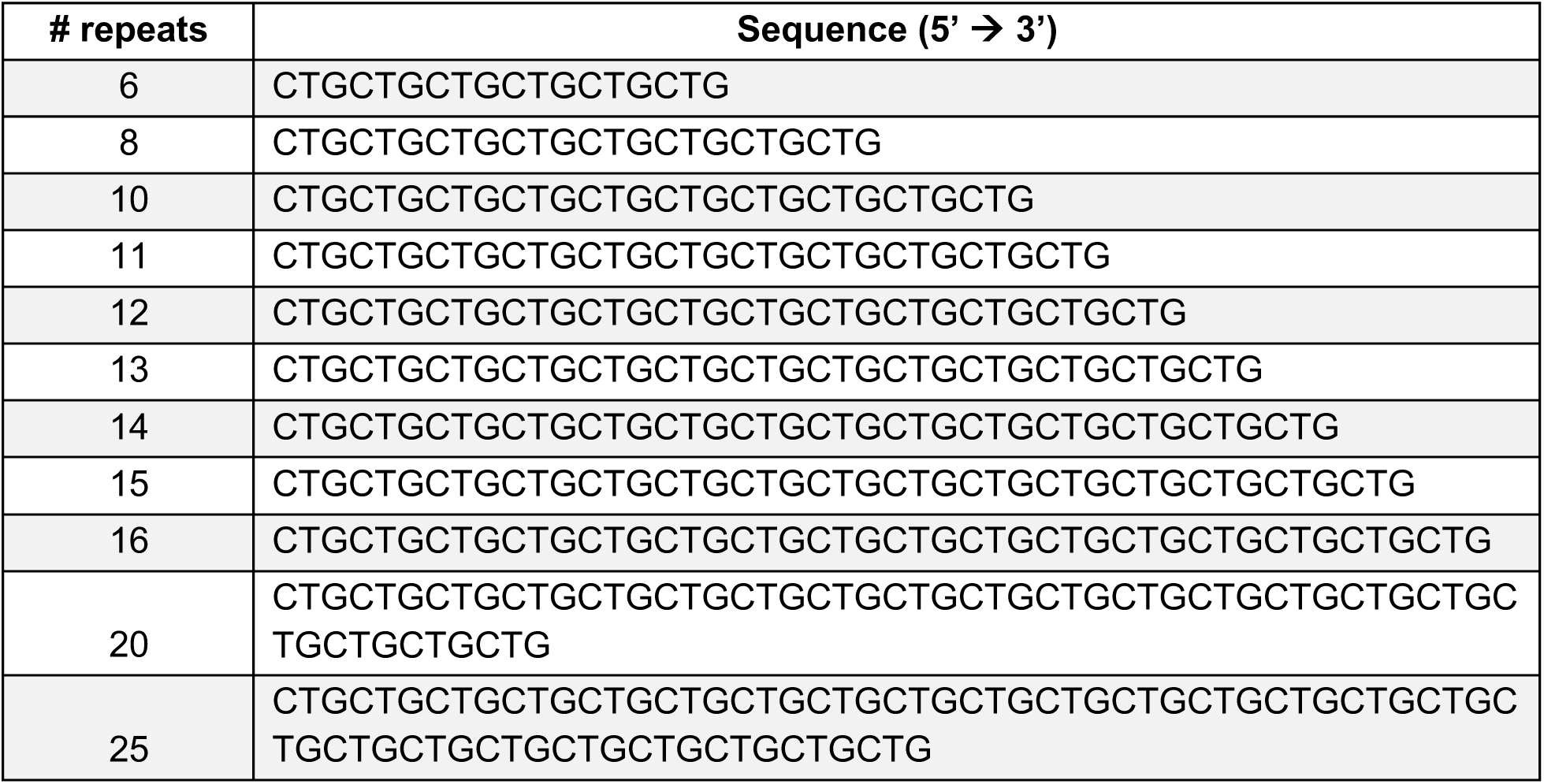
CTG_n_ repeats for melting profile analysis in Fig. 3A.

**Supplementary Table 14.**
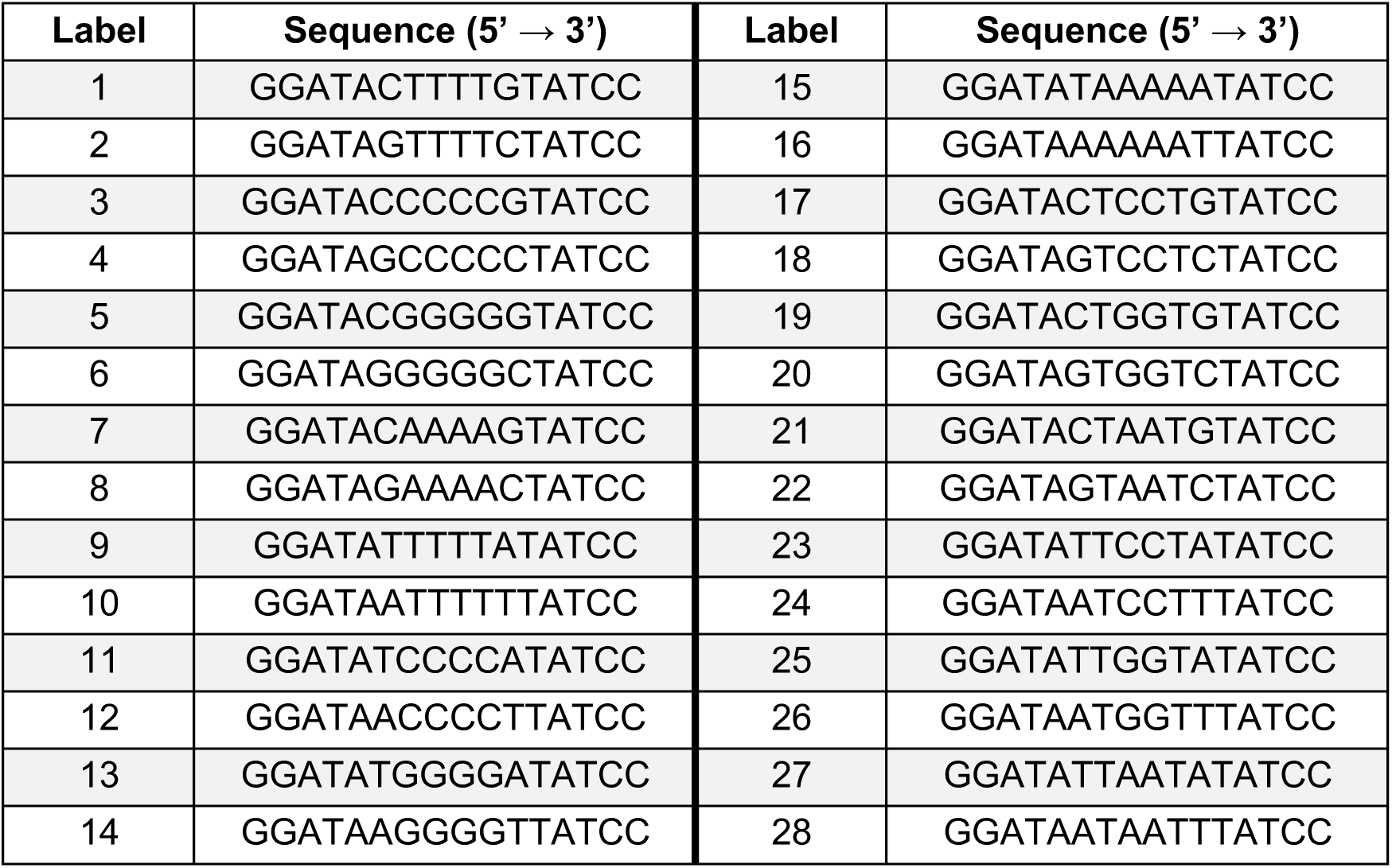
Hairpin sequences for melting profile analysis in Fig. 4A.

**Supplementary Table 15.**
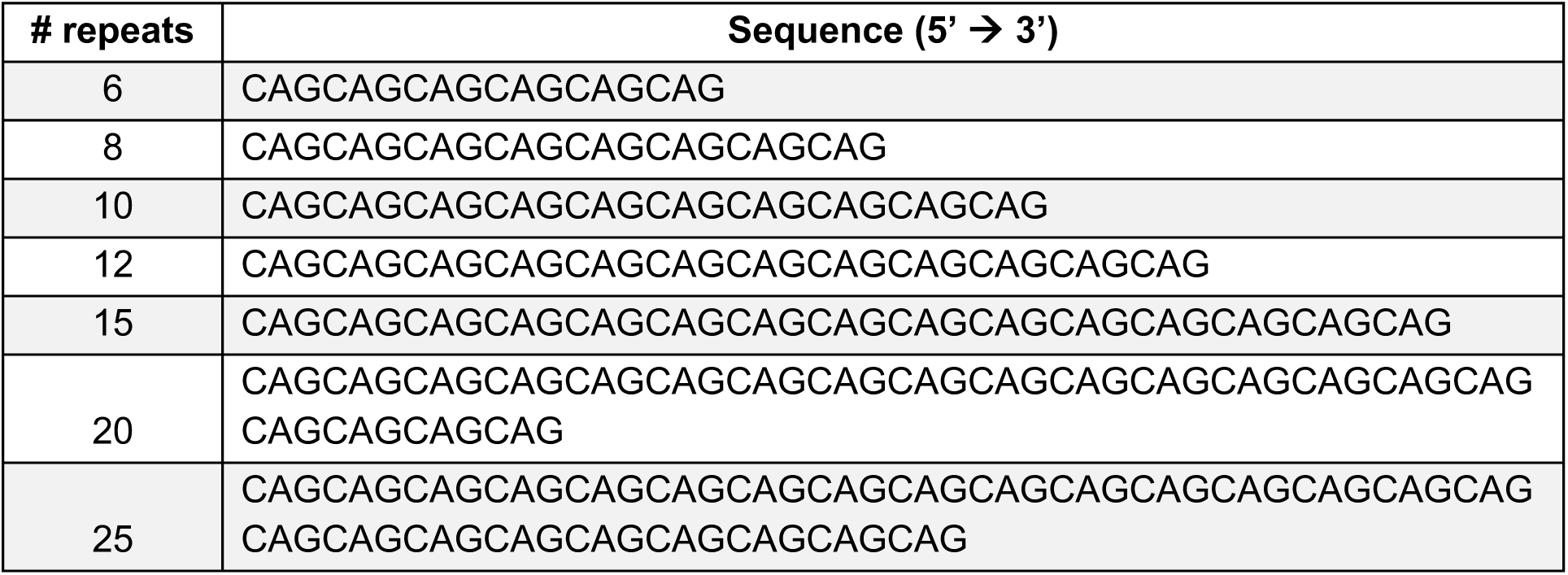
CAG_n_ repeats for melting profile analysis in Supplementary Fig. 1.

**Supplementary Table 16.**
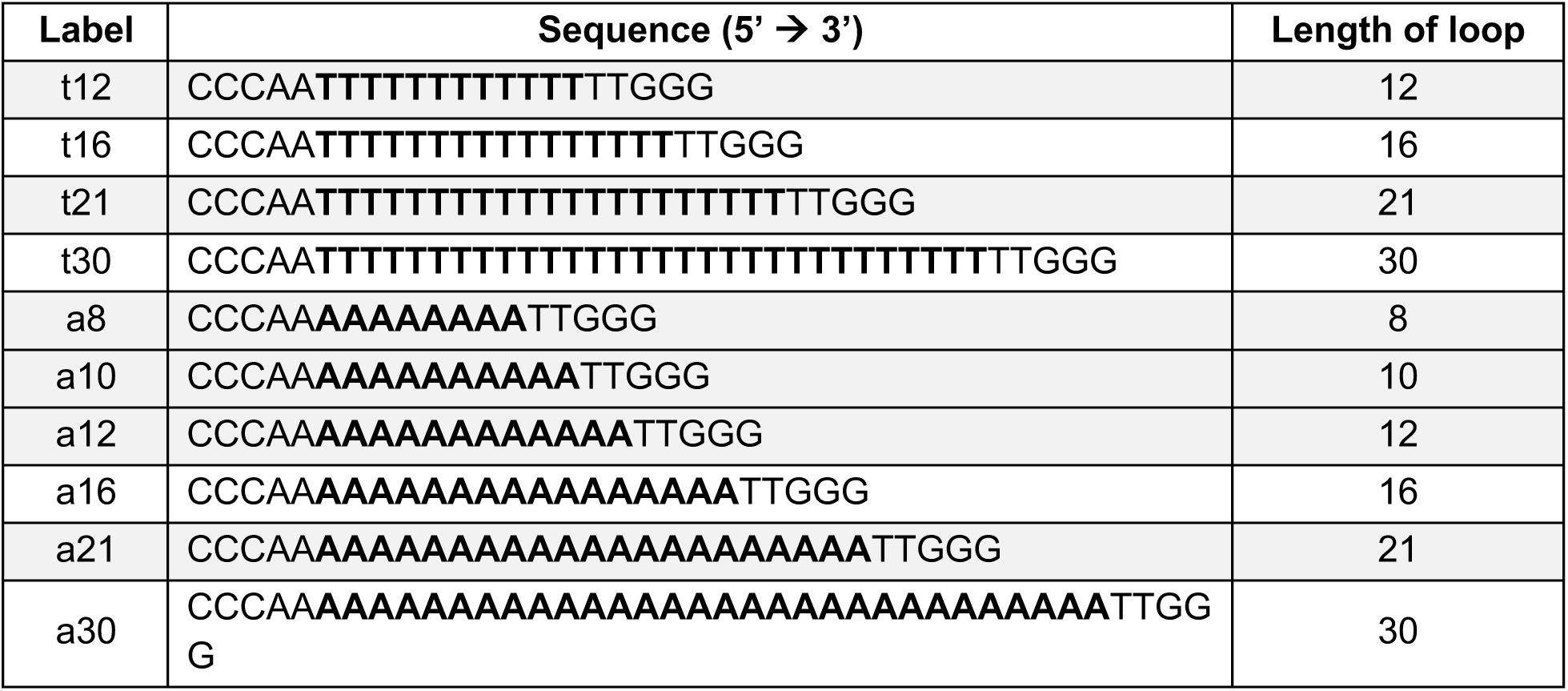
Hairpin melting profile analysis in Fig. 4B, and the loop sequence is bolded.

**Supplementary Table 17.**
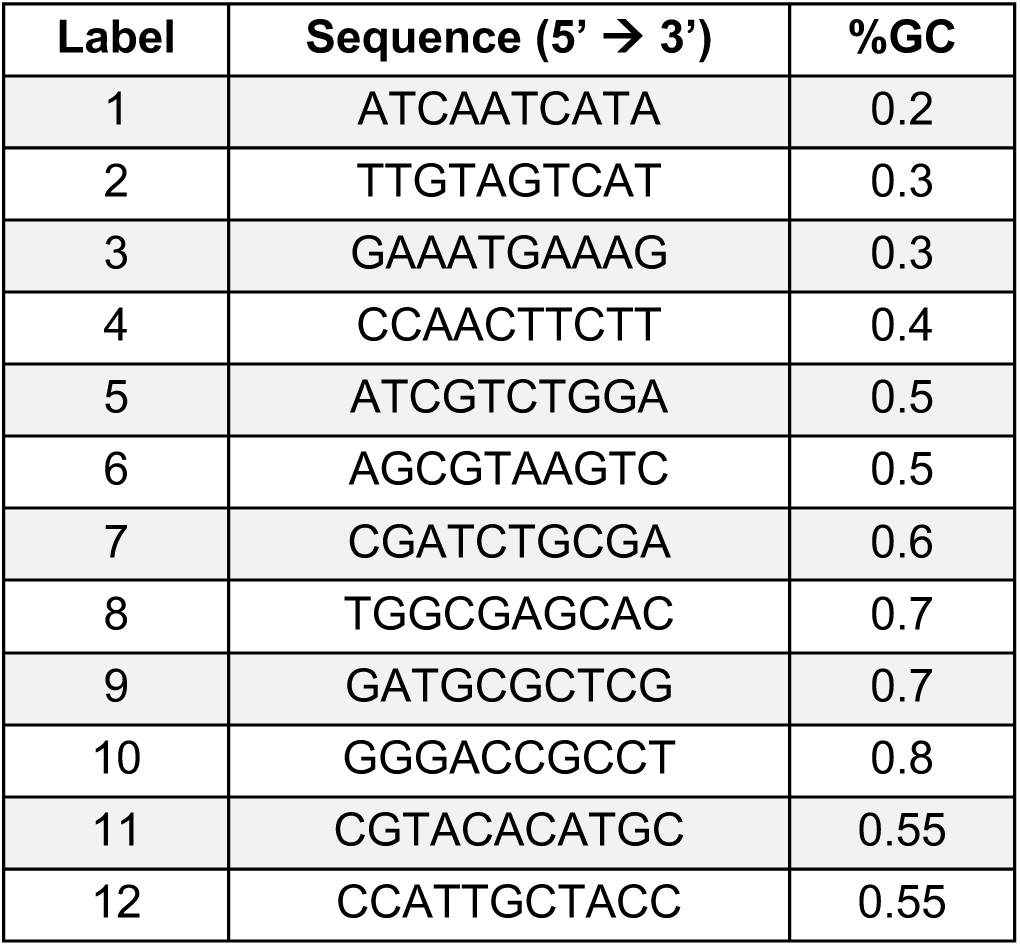
Duplex melting profile analysis in Fig. 4B.

**Supplementary Table 18.**
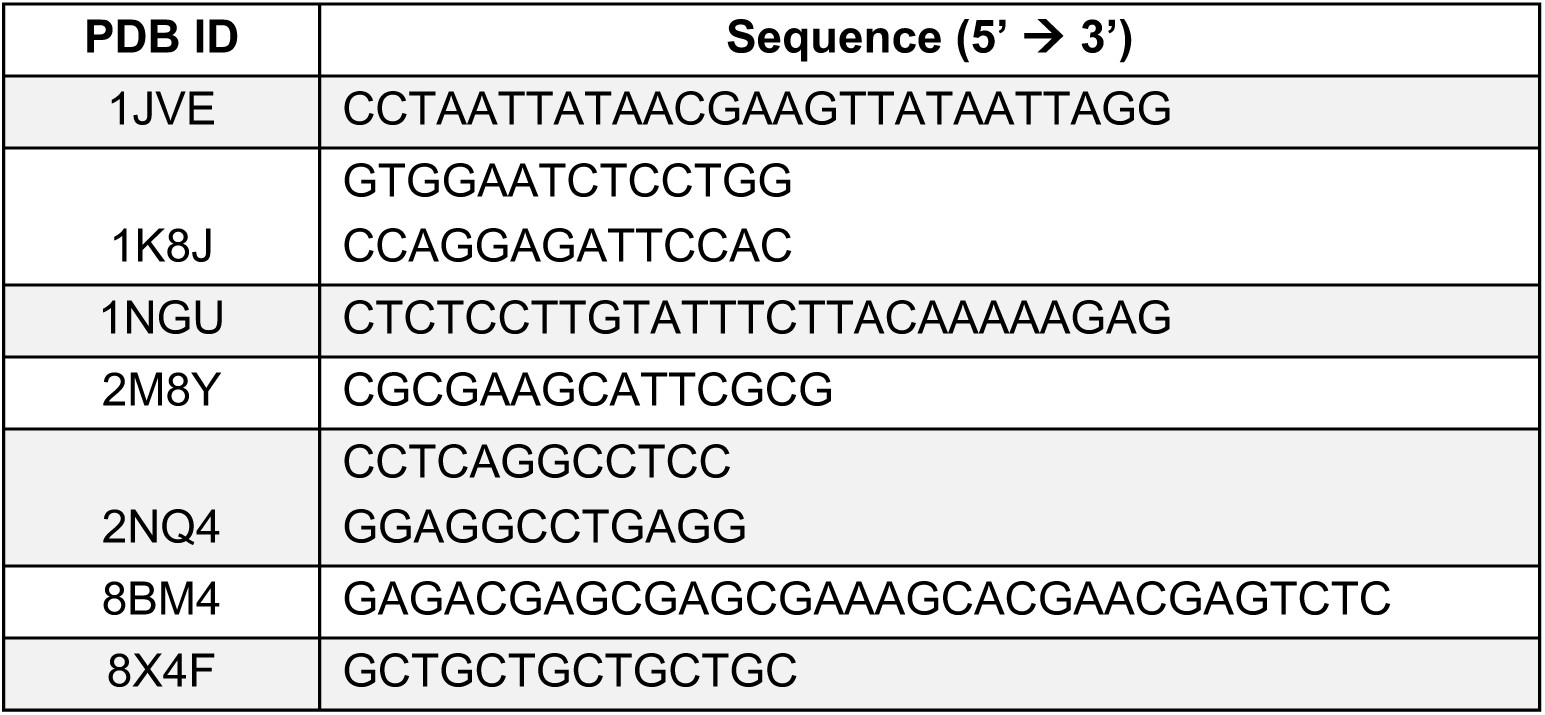
PDB and sequence used for folding simulation in Fig. 4C.

**Supplementary Figure 1.**
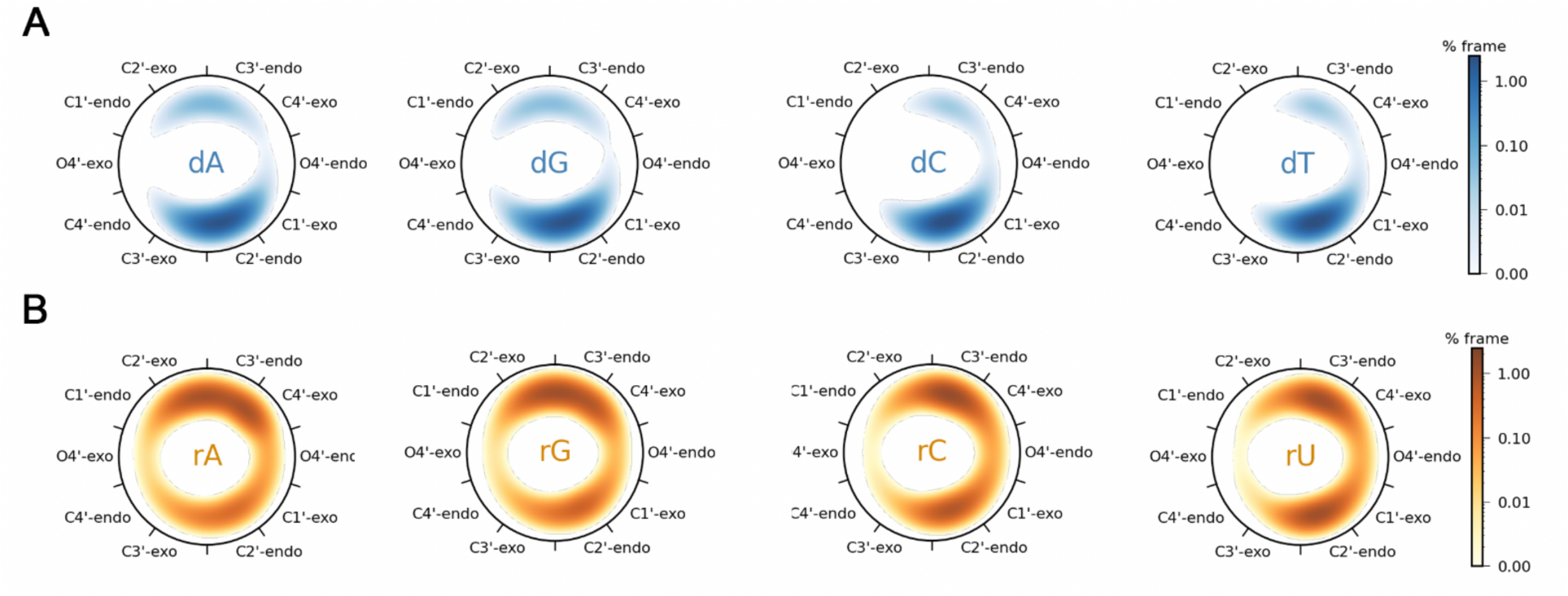
Comparison of sugar phase pseudorotation between (A) DNA and (B) RNA. Histograms mapped to a 2D coordinate plotting the pseudorotation phase angle, calculated via the Cremer and Pople (CP) method, with a bin of one degree, and smoothed with a periodic Gaussian filter (σ = 5°). The color represents the population intensity with a logarithmic scale, calculated as percentage of total frames per degree.

**Supplementary Figure 2.**
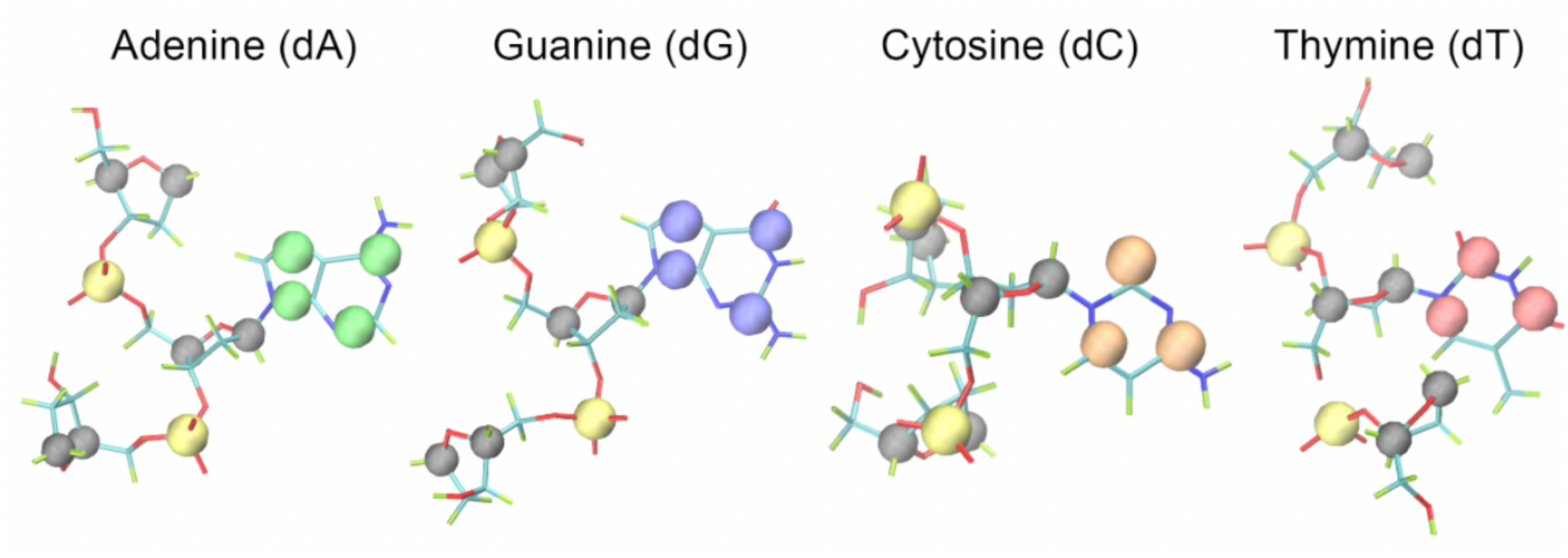
All-atom structure of abasic triphosphate overlaid with CG beads. The abasic tri-nucleoside molecule consists of three deoxyribose sugar rings, two phosphate groups, and one nucleobase, and is used to generate the reference atomistic simulation for bonded term parameterization. The corresponding CG beads are colored in black for sugar, yellow for phosphate group, and different colors for each nucleobase. Adenine is colored in green, guanine in blue, thymine in red, and cytosine in orange. The physical volumes of the CG beads are not drawn to scale in this schematic.

**Supplementary Figure 3.**
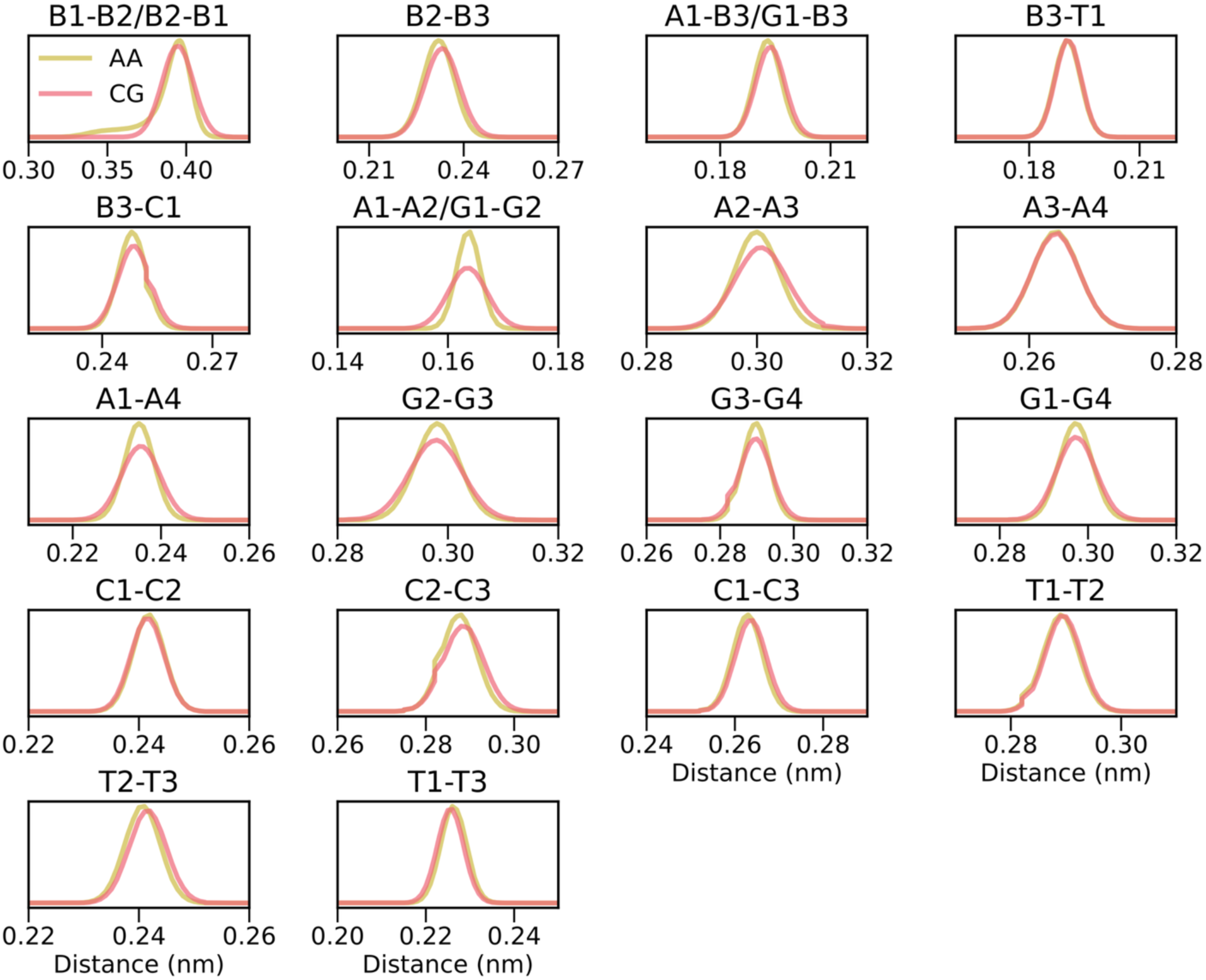
All-atom and CG distribution for bond. The distribution between all-atom (yellow) and corresponding CG (red).

**Supplementary Figure 4.**
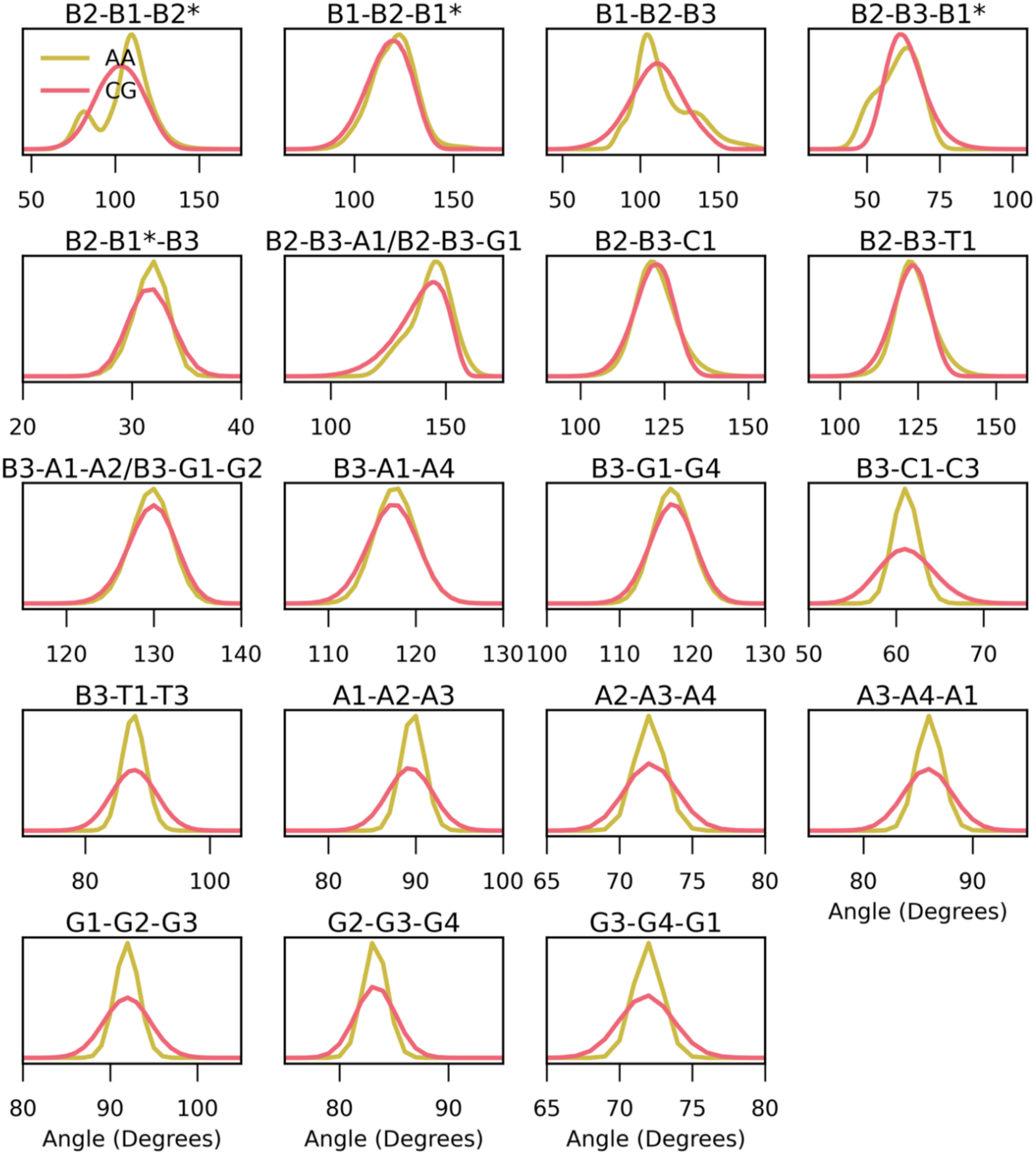
All-atom and CG distribution for angle. The distribution between all-atom (yellow) and corresponding CG (red). *Represents the bead in the next residue.

**Supplementary Figure 5.**
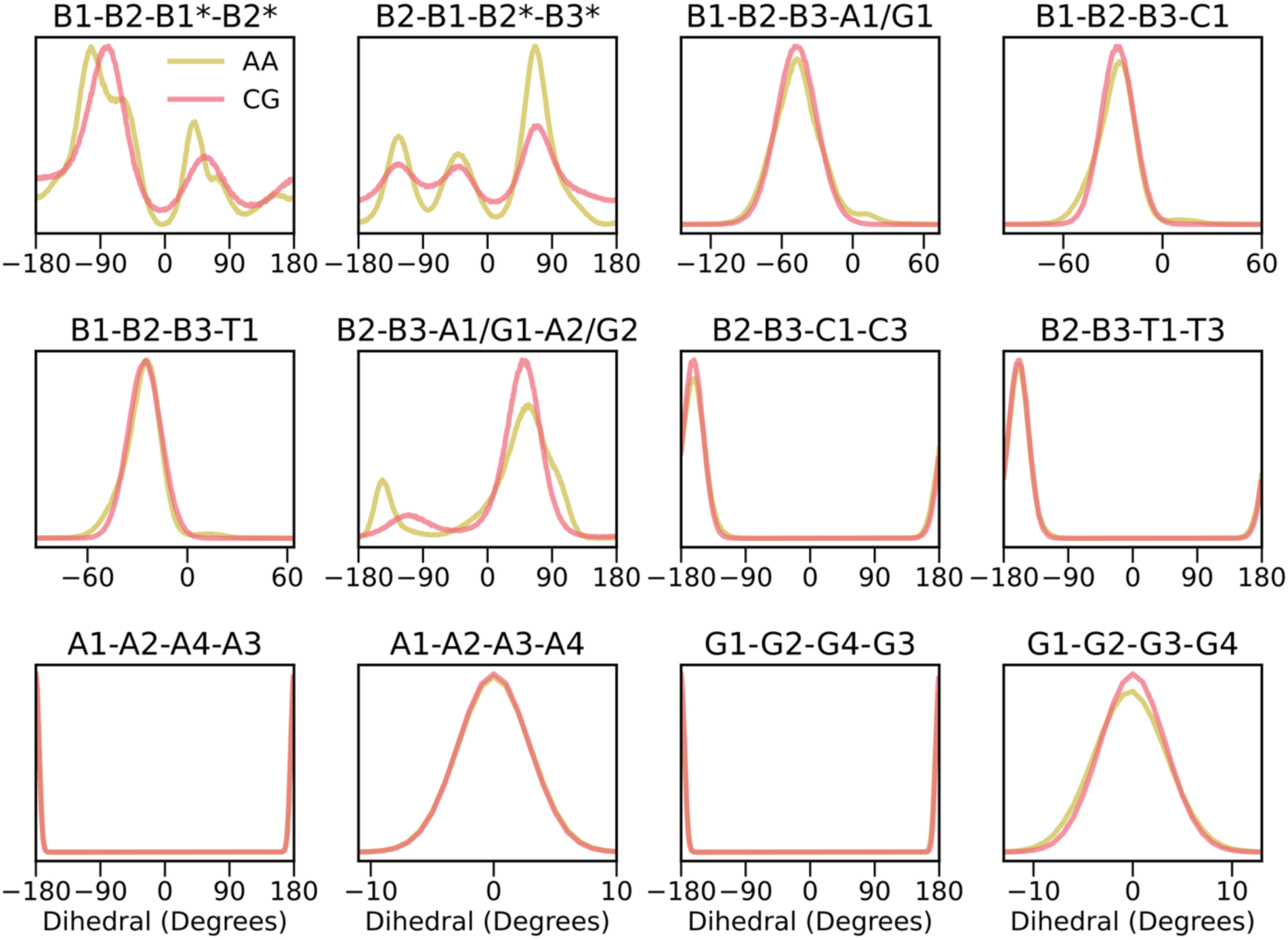
All-atom and CG distribution for dihedral. The distribution between all-atom (yellow) and corresponding CG (red). * Represents the bead in the next residue.

**Supplementary Figure 6.**
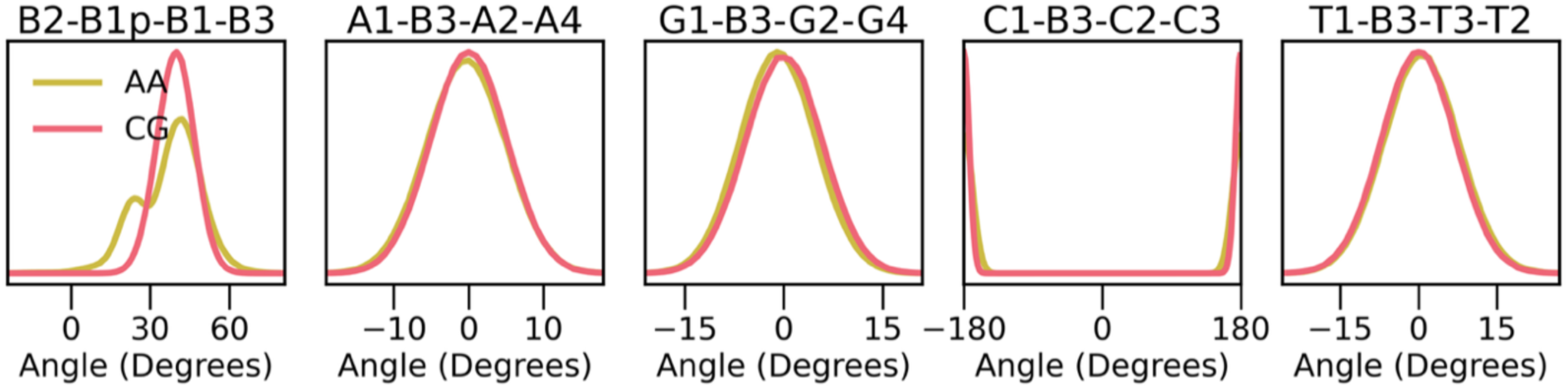
All-atom and CG distribution for improper. The distribution between all-atom (yellow) and corresponding CG (red). * Represents the bead in the next residue.

**Supplementary Figure 7.**
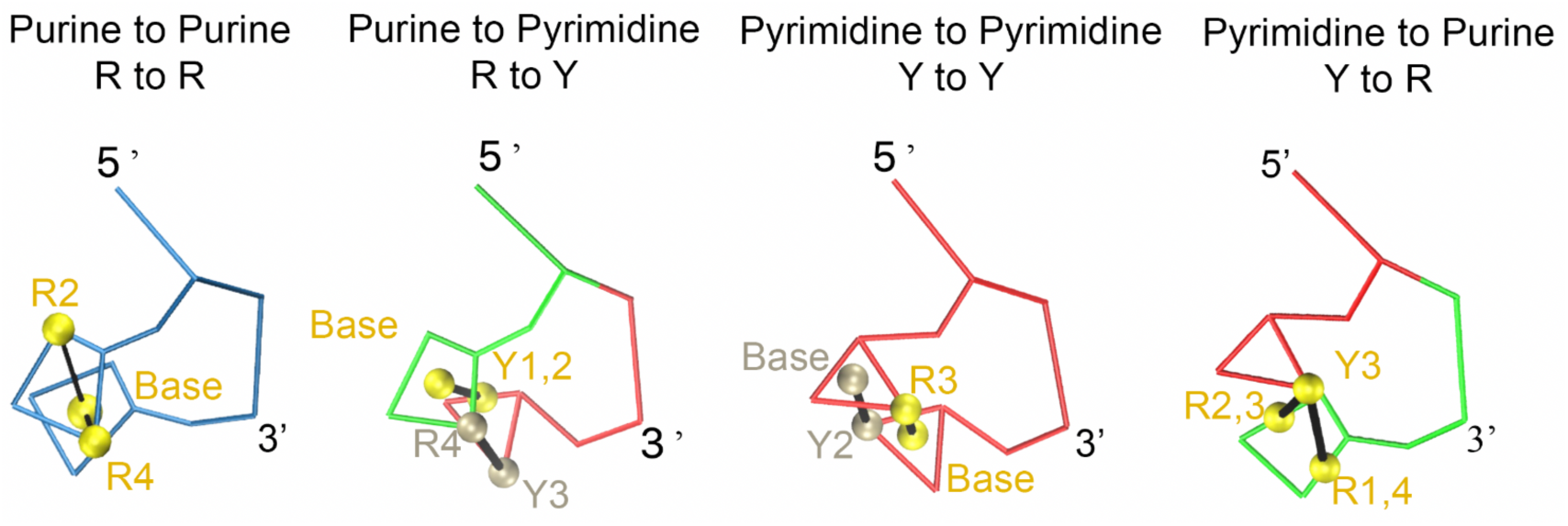
Base stacking design. Base stacking is assigned to consecutive 5’ to 3’ neighbors via two virtual bonds (drawn in black) and virtual sites (colored in yellow/silver). The stacking schemes are classified by purine (R: A/G) or pyrimidine (Y: C/T) pairs (RR, RY, YY, YR). The beads that are used to calculate the center of geometry for the virtual site are listed in yellow or silver text, and the label “Base” refers to the center of geometry of the nucleobase. Individual nucleotide is color coded with green, blue, red, and orange, indicating adenine, guanine, thymine, and cytosine, respectively. Further details for each stacking design are listed in **Supplementary Table 11**.

**Supplementary Figure 8.**
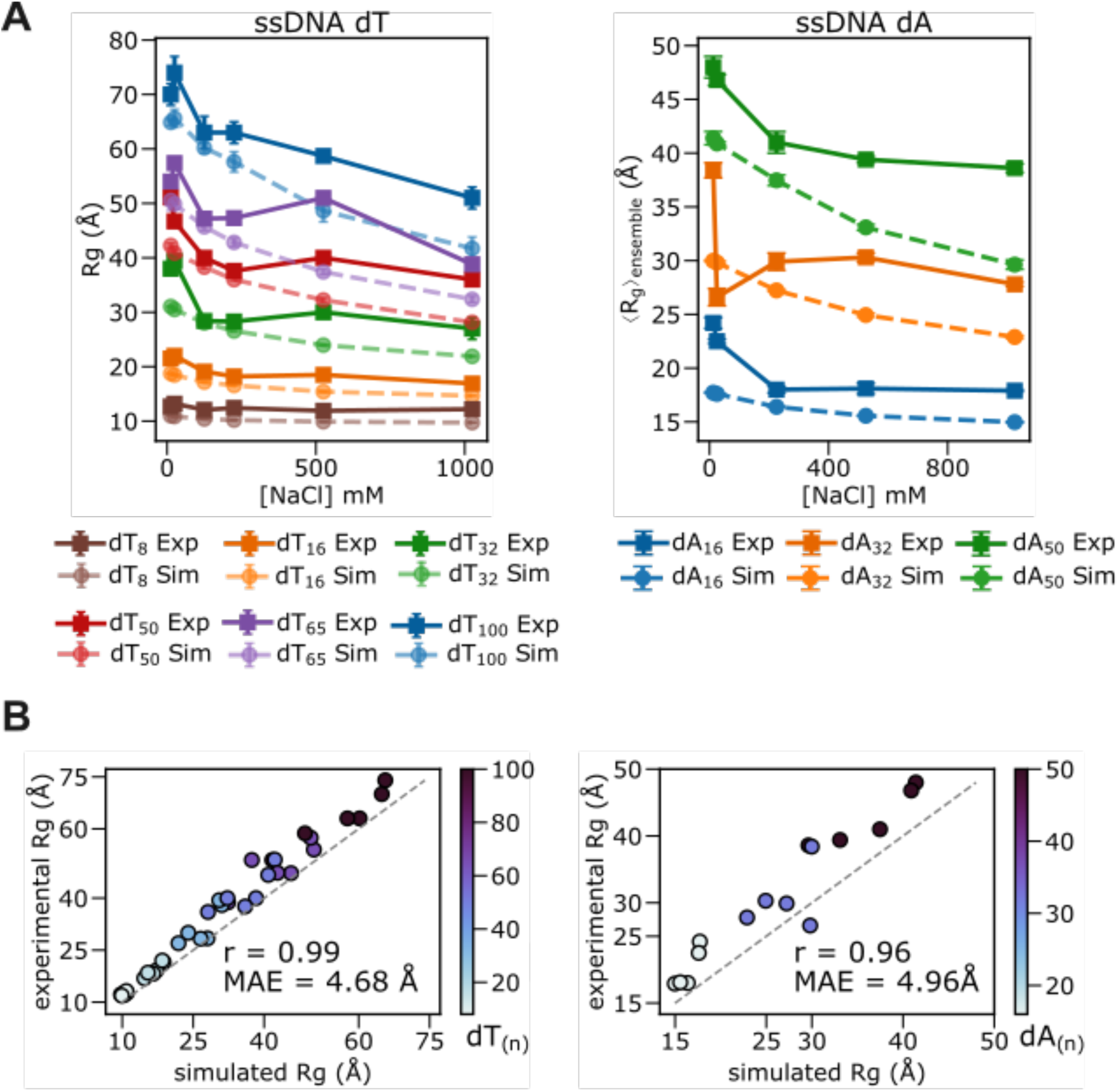
Agreement between simulated and experimentally measured R_g_ for poly-dT (left) and poly-dA (right). **(A)** R_g_ as a function of salt (NaCl) in different numbers of bases for ssDNA dT and dA. The simulation data are shown in dash, and experimental data as a solid line. **(B)** Pearson correlation between simulated and experimental R_g_ for poly-dT and poly-dA. Pearson correlation coefficient (r) and mean absolute error (MAE) are displayed in the graph.

**Supplementary Figure 9.**
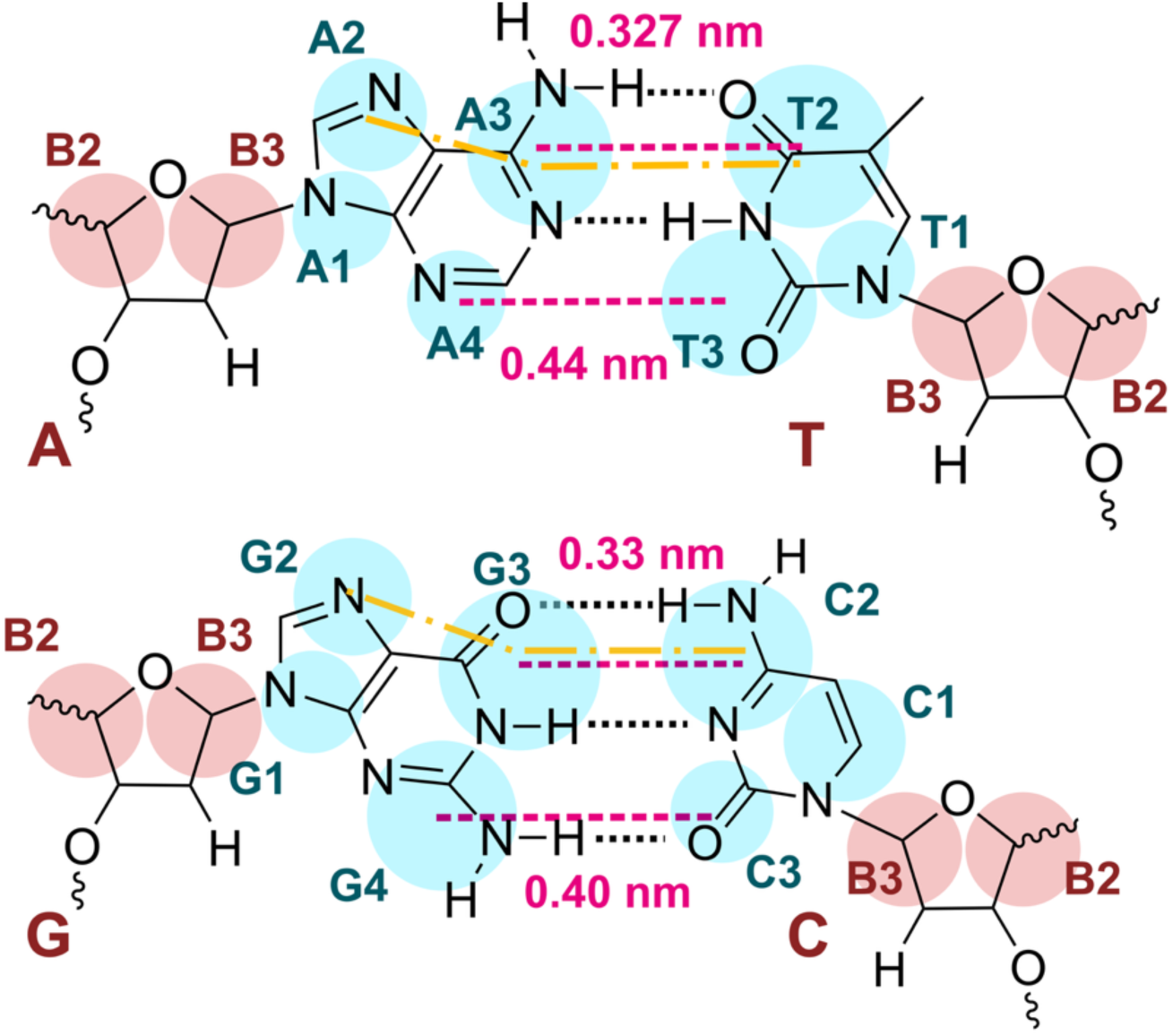
The beads involved in base pairing for A-T and G-C. Base beads are colored in cyan, and sugar beads are colored in red. The atomistic-resolution hydrogen bond is indicated by dash black line. The base-pairing participating beads are connected by a pink dashed line, and the equilibrium distance is listed in pink text. The beads used for angle calculation are connected by a yellow dashed line. The equilibrium distances and angles are listed in **Supplementary Tables 8** and **9**, respectively.

**Supplementary Figure 10.**
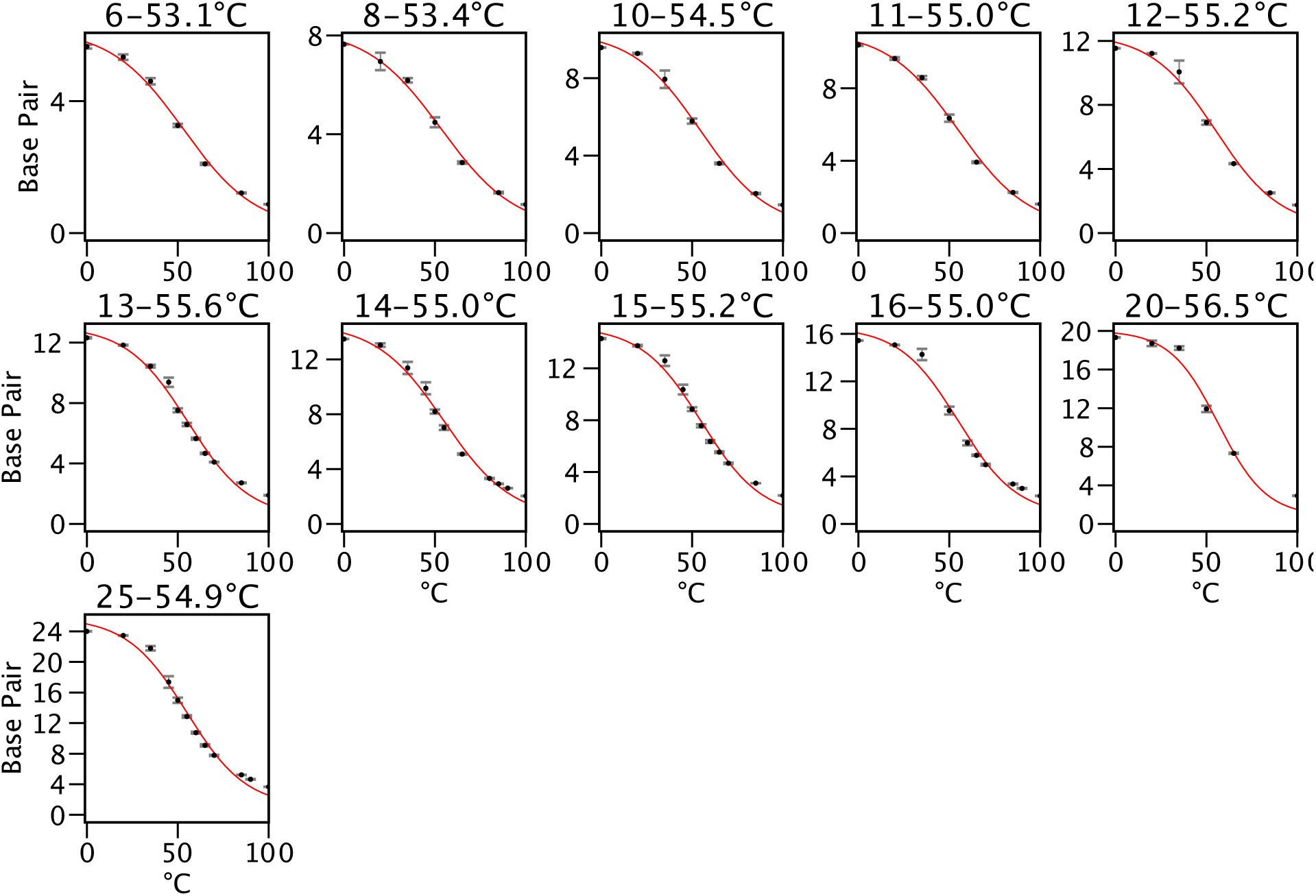
The melting profiles of CTG_n_ in Fig. 4A. The title shows the label of the hairpin and the melting temperature. The corresponding sequences are listed in **Supplementary Table 13**. Each data point represents the block average from the CG 6 µs simulation, with the first 50ns discarded.

**Supplementary Figure 11.**
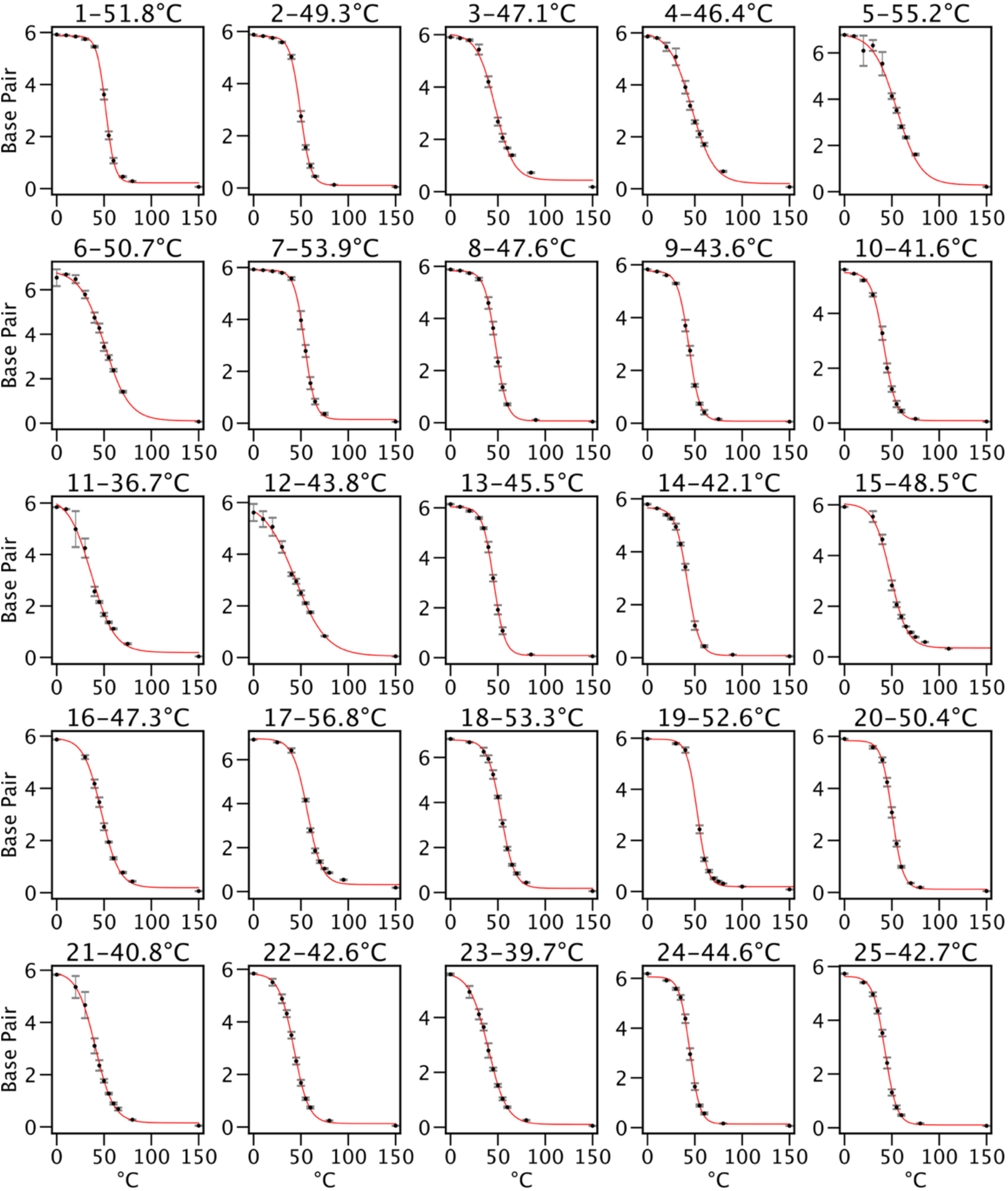
The melting profiles of hairpins in Fig. 4A. The title shows the label of the hairpin and the melting temperature. The corresponding sequences are listed in **Supplementary Table 14**. Each data point represents the block average from the CG 6 µs simulation, with the first 50ns discarded.

**Supplementary Figure 12.**
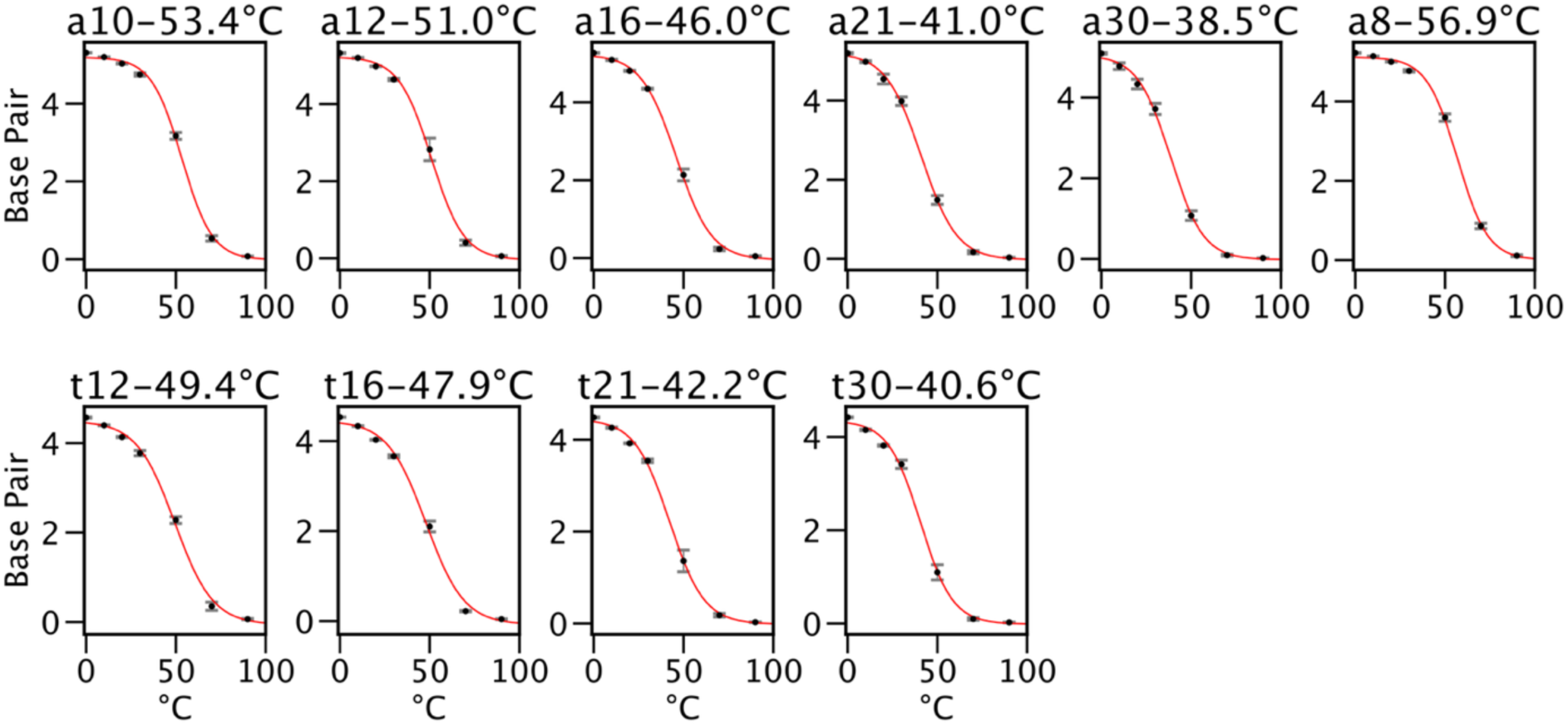
The melting profiles of the hairpin in Fig. 4B. The title shows the label of the hairpin and the melting temperature. The corresponding sequences are listed in **Supplementary Table 16**. Each data point represents the block average from the CG 6 µs simulation, with the first 50ns discarded.

**Supplementary Figure 13.**
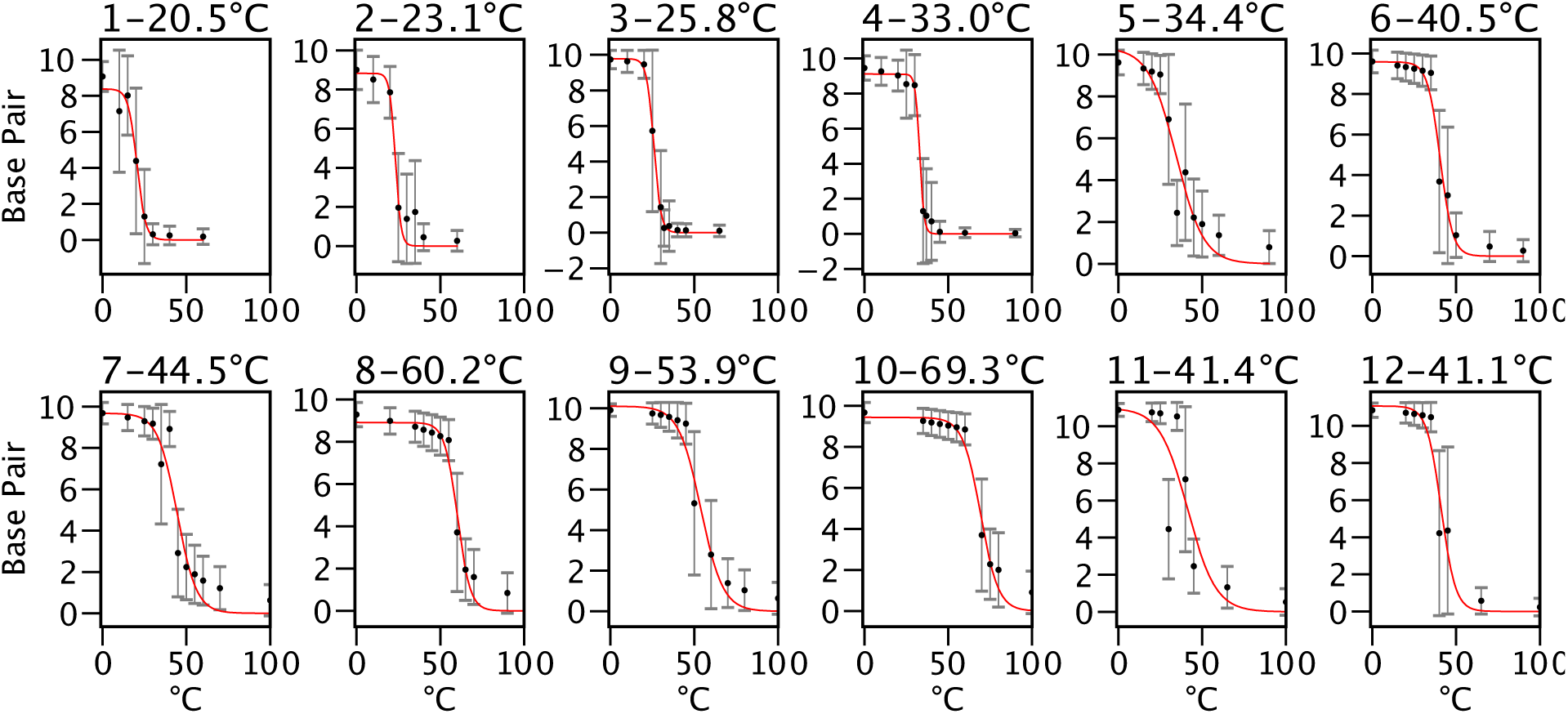
The melting profiles of duplexes from **Fig. 4B**. The title shows the label of the duplex and the melting temperature. The corresponding sequences are listed in Supplementary Table 12. Each data point represents the block average from the CG 10 µs simulation, with the first 200ns discarded.

**Supplementary Figure 14.**
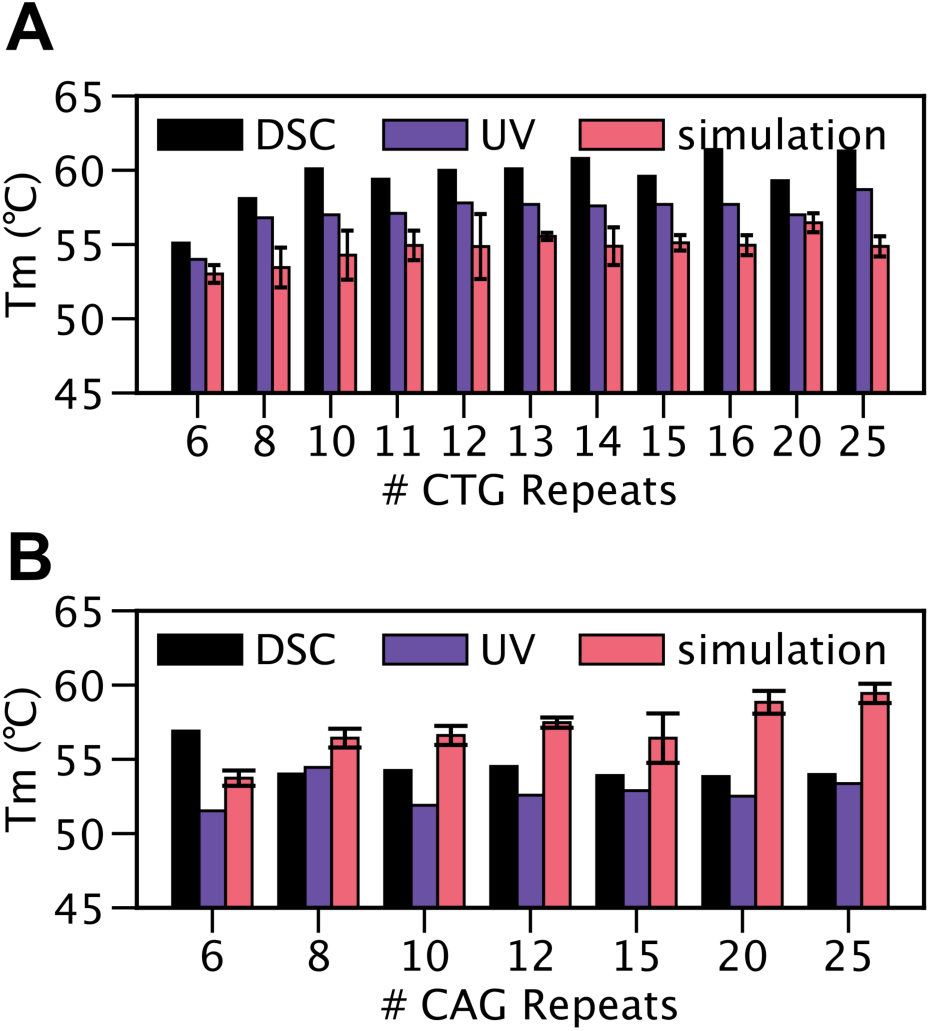
Compare experimental and simulated melting profiles of hairpin. (**A**) CTG_n_ repeats and (**B**) CAG_n_ repeats. The error bar represents the fitted melting temperature from the block average of the CG 6 µs simulation, with the first 50ns discarded.

**Supplementary Figure 15.**
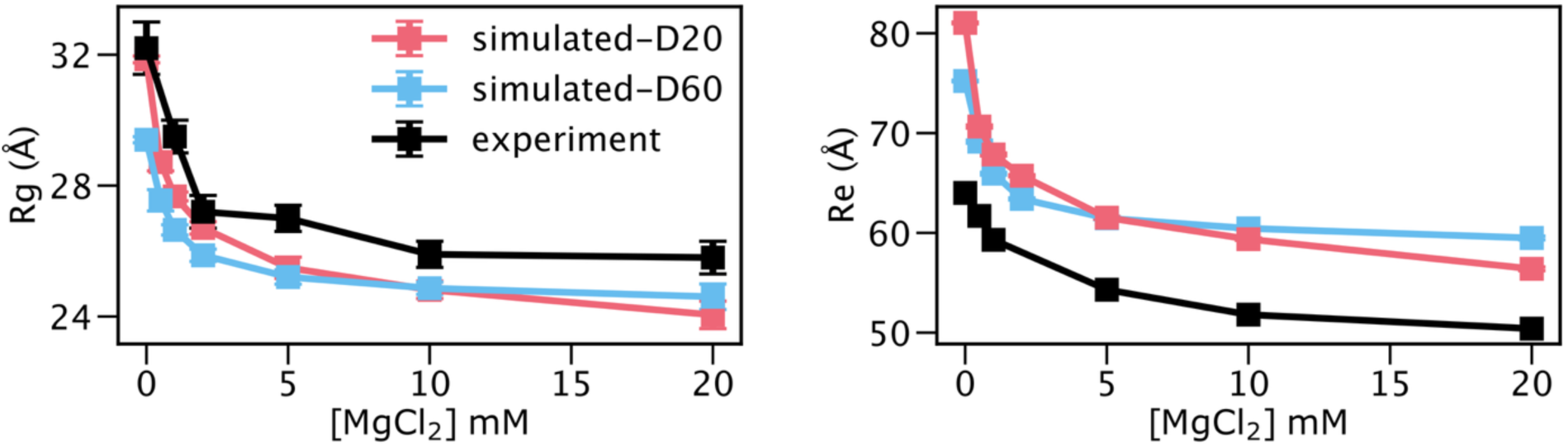
Effect of dielectric constant on ssDNA properties. dT_30_ R_g_ and R_e_ under different MgCl_2_ concentrations simulated with dielectric constants of 20 (pink) and 60 (blue) compared to experiment (black).

